# Cellular metabolic state controls mitochondrial RNA kinetics

**DOI:** 10.1101/2025.05.13.653903

**Authors:** Sean D. Reardon, Clarisa Bautista, Shannon C. Cole, Julien Cicero, Tatiana V. Mishanina

## Abstract

Human mitochondrial genome encodes essential genes for the oxidative phosphorylation (OXPHOS) complexes. These genes must be transcribed and translated in coordination with nuclear-encoded OXPHOS components to ensure correct stoichiometry during OXPHOS complex assembly in the mitochondria. While much is known about nuclear gene regulation during metabolic stresses like glucose deprivation, little is known about the accompanying transcriptional response in mitochondria. Using microscopy, roadblocking qPCR, and transcriptomics, we studied mitochondrial transcription in cells subjected to glucose deprivation, which is known to cause nuclear transcription downregulation and to activate the integrated stress response (ISR). We found that glucose deprivation stabilizes mitochondrial RNAs and slows mitochondrial transcription, effects that are quickly reversed with glucose reintroduction. Although transcriptomics revealed strong upregulation of the ISR, mitochondrial RNA stabilization was not upregulated by pharmacological activation of the ISR, but was promoted by inhibition of glycolysis, unveiling a direct connection between metabolism and regulation of mitochondrial gene expression.

## Introduction

The mitochondria in eukaryotic organisms contribute the majority of cellular ATP through OXPHOS. Mitochondria contain multiple copies of their own genome and their own ribosomes, which must coordinate with the nuclear genome and cytosolic ribosomes to ensure the correct stoichiometry of the OXPHOS complex proteins synthesized in the cytosol and mitochondrial matrix^1,2^. Incorrect stoichiometry of OXPHOS proteins results in the activation of the unfolded protein response (UPR) and cell sensitivity to proteosome inhibition^1^. Thus, coordination between the nuclear and mitochondrial genomes is essential to cellular health. The human mitochondrial genome (mtDNA) is a ~16 kb circular DNA encoding 13 essential OXPHOS complex proteins, 22 tRNAs, and 2 rRNAs (Fig. 1A). The mtDNA is entirely maintained, transcribed, and replicated by nuclear-encoded proteins^3^, with transcription of human mtDNA carried out by the mitochondrial RNA polymerase (POLRMT) and the transcription factors: Mitochondrial Transcription Factor A (TFAM), Mitochondrial Transcription Factor 2B (TFB2M), and the Mitochondrial Transcription Elongation Factor (TEFM)^4–6^. Of these factors, TFAM also compacts individual mtDNA molecules into nucleoids, and TFAM concentration is thought to affect the accessibility of mtDNA for processes like transcription and replication^7–10^.

**Figure 1.**
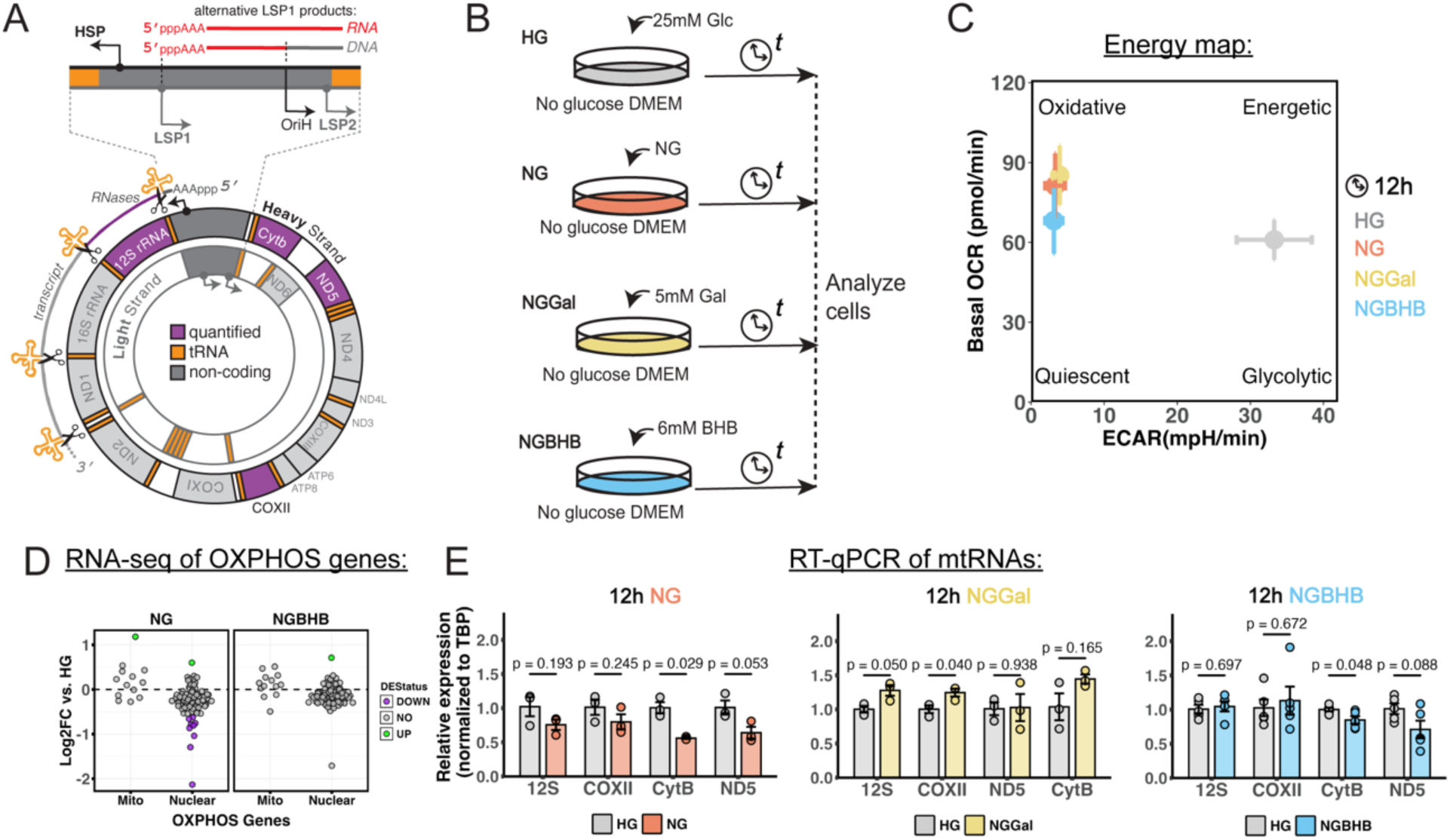
Media manipulations in this study and their consequences to cellular processes. **A)** Depiction of the human mitochondrial DNA with genes encoded by the heavy strand shown on the outside and genes encoded by the light strand on the inside. Mitochondrial genes quantified throughout this study are highlighted in purple. **B)** Schematic of color-coded cell treatments used throughout the study. **C)** Energy map of HEK293T cells deprived of glucose for 12 h showing the shift to oxidative metabolism. Data are from a Seahorse extracellular flux assay, with 10 wells per condition. **D)** RNA-seq based log_2_ fold-change in expression of nuclear and mitochondrially encoded OXPHOS proteins after a 12 h NG or NGBHB treatment, compared to HG. Points represent individual genes with green as upregulated, grey as non-significant, and purple as downregulated (n = 3 biological replicates). **E)** Quantitative reverse transcription PCR (RT-qPCR) analysis of representative mitochondrial transcripts shows steady pools of these RNAs after 12 h of glucose deprivation (n = 3-5 biological replicates, normalized to tata-box-binding protein (TBP), data are means ± S.E.M. and associated *p* values).

Mitochondrial RNA (mtRNA) is synthesized as a polycistronic transcript from 3 promoters, the heavy strand promoter (HSP) and light strand promoters 1 and 2 (LSP1 and LSP2), located within the non-coding region of mtDNA (Fig. 1A). Recent studies demonstrated that mitochondrial mRNA transcripts are ~100-fold more abundant than nuclear OXPHOS-encoding mRNAs, are rapidly processed co-transcriptionally, and found as residents of a cellular body separate from mtDNA known as the mitochondrial RNA granule (MRG)^11,12^. mtRNA is both synthesized and degraded much more quickly than nuclear RNA, with mtRNA stability mainly determined by the Leucine-Rich Pentatricopeptide Repeat Containing protein (LRPPRC)^12^. Both mtRNA synthesis and degradation are ATP-dependent^6,13^, making rapid mtRNA kinetics an energy-expensive process for the cell. Finally, mtDNA replication is dependent on mitochondrial transcription where the terminated RNA transcript from LSP1 primes replication of the heavy mtDNA strand (Fig. 1A), although it is not known if full-length mtDNA transcription and replication are mutually exclusive on the same mtDNA nucleoid or what factors may regulate these processes^14,15^.

Major progress has been made in classifying the machinery and kinetics that dictate mitochondrial transcription and replication in standard cell culture conditions of abundant glucose. However, questions remain about how metabolic changes, like a shift from a glycolytic to an oxidative metabolism, or metabolic stress like glucose deprivation, may impact mitochondrial transcription and replication processes in the short term.

Here, we sought to determine short-term (4-12 hours, h) effects of metabolic stress on mitochondrial transcription and mtDNA replication in cells cultured under glucose deprivation, with and without supplementation of alternative carbon sources. Specifically, we compared cells grown in Dulbecco’s Modified Eagle Medium (DMEM) without an extra carbon source; substituting glucose with galactose, a traditional way to test OXPHOS function^16^, or substituting glucose with β-hydroxybutyrate (BHB), the most abundant ketone body in mammals used as an alternative fuel during fasting, prolonged exercise, or when subject to a ketogenic diet (Fig. 1B; NG, NGGal, and NGBHB, respectively)^17,18^. Galactose can enter glycolysis through the Lenior pathway but is considered a poor glycolytic substrate, leading cells to shift to OXPHOS^19^ (Fig. 1C). BHB is a four-carbon short-chain fatty acid produced predominantly in the liver when glucose becomes scarce in mammals. BHB circulates through the blood to other organs, most notably the brain, to support their continued function through oxidative metabolism^18^. As a fuel, BHB enters cells and is first broken down within mitochondria by BHB dehydrogenase to acetoacetate, then to acetoacetyl-CoA by 3-ketoacyl-CoA transferase and finally into two acetyl-CoA molecules, which can enter citric acid cycle to supply reducing equivalents for OXPHOS. Recent work showed that BHB indeed increases ATP levels in neurons, suggesting that it is directly used as a carbon source^20^. All three conditions shift energy production from glycolysis to OXPHOS to maintain constant cellular concentration of ATP ([ATP]) short-term (Fig. 1C and S1A).

Importantly, a lack of glucose causes unfolded proteins to accumulate in the endoplasmic reticulum (ER), triggering ER stress and the integrated stress response (ISR), a cellular program facilitating several adaptations to conserve ATP and keep cells alive including slowed global translation, upregulation of OXPHOS, and selective transcription of genes aimed at maintaining cellular energy homeostasis and protein folding^16,21,22^. The ISR utilizes several key transcription factors (ATF4, DDIT3, and ATF6B), all of which are upregulated at the transcript level in our conditions as evident from the RNA sequencing (RNA-seq) analysis of 12 h NG and NGBHB treated samples. Additionally, Gene Ontology (GO) term enrichment of differentially expressed genes shows upregulation of ER stress-related genes and downregulation of genes related to cell proliferation (Fig. S1B-S1E). These results are complemented by the halt in cell proliferation in our conditions but maintained cell viability, indicative of an adaptive ISR (Fig. S1F and S1G).

Using the above glucose deprivation conditions, with a backdrop of an energetic crisis and adaptive ISR, we provide a timeline for fine-tuned mitochondrial gene expression. We show that cellular energetic status and specifically decreased mitochondrial [ATP] coincides with a slowdown in mtRNA synthesis and degradation, but that a steady-state pool of mtRNA is maintained. Strikingly, this effect is glucose-dependent and fully reversible upon re-introduction of glucose, confirming a long-held hypothesis that mitochondrial transcription senses the metabolic status of the cell. We find that the slowdown in mtRNA degradation and synthesis observed under glucose deprivation is reproduced in nutrient abundance by direct inhibition of glycolysis but not pharmacological induction of ISR signaling. Finally, we show that the slowdown in mtRNA degradation and mitochondrial transcription is largely independent of changes in mtDNA replication, with the exception of the combined glucose deprivation and BHB supplementation where mtDNA replication was stimulated only in HEK293T cells. Our study provides evidence that mitochondrial transcription and mtRNA degradation processes sense the cellular energetic status to tune the lives of mitochondrial transcripts in the face of metabolic stress.

## Results

### Glucose deprivation decreases nuclear-encoded OXPHOS subunit transcripts and stabilizes mature mitochondrial RNAs

In light of the need to conserve energy under glucose deprivation and the ~100-fold mismatch between nuclear vs mitochondrial OXPHOS subunit mRNA expression, we expected to see drastic changes in mitochondrial gene expression under glucose deprivation. Fold-change analysis from RNA-seq of nuclear-encoded OXPHOS subunits showed a general downregulation in both NG and NGBHB conditions compared to HG, with a stronger effect in NG (Fig. 1D). However, fold-changes for mitochondrially encoded OXPHOS genes did not show such a general downregulation (Fig. 1D), consistent with other studies using substitution of glucose with galactose^16,23^. We confirmed the unchanged steady-state pools of mtDNA-encoded transcripts, using whole-cell RNA-seq analysis and reverse-transcription quantitative PCR (RT-qPCR) of representative mtRNAs (purple genes in Fig. 1A and data in Fig. 1F) against the TATA-box Binding Protein (TBP). This observation is consistent with previously published studies that treated cells with galactose^16,23^. Cells thus maintain a steady pool of mtRNAs during glucose deprivation. Given the energetic burden to sustain large excess of mitochondrial transcripts over the nuclear OXPHOS transcripts (100:1, Fig. S1H), we chose to analyze mtRNA kinetics (i.e., RNA synthesis vs degradation) as a possible regulatory point.

To probe the kinetics of the steady-state pools of mtRNA, we deployed a recently reported roadblocking qPCR strategy to measure relative turnover of individual mtRNA transcripts between media conditions^24^. With this technique, the newly synthesized RNA is metabolically labelled with the uridine analog 4-thiouridine (4sU), whereas the unlabeled RNA reports on the pre-existing transcripts prior to labelling (Fig. 2A). Cells are treated with 4sU for a given time, and total RNA is purified and treated with *N*-ethylmaleimide (NEM) to alkylate the sulfhydryl group on the 4sU incorporated into RNA. These alkylated bases interrupt complementary DNA (cDNA) synthesis during the reverse transcription step of RT-qPCR sample preparation, while full-length cDNA of unlabeled transcripts is synthesized. The resulting cDNA is then used to compare the relative unlabeled RNA between conditions normalized by a synthetic spike-in RNA control. The qPCR signal allows relative comparisons of RNA turnover between conditions or timepoints, where the signal reports on the unlabeled (older) RNA of interest (Fig. 2B). Using this technique, we chose to label for the last 2 h of each treatment using a final concentration of 50 µM 4sU, a concentration known to limit 4sU toxic effects^25^.

**Figure 2.**
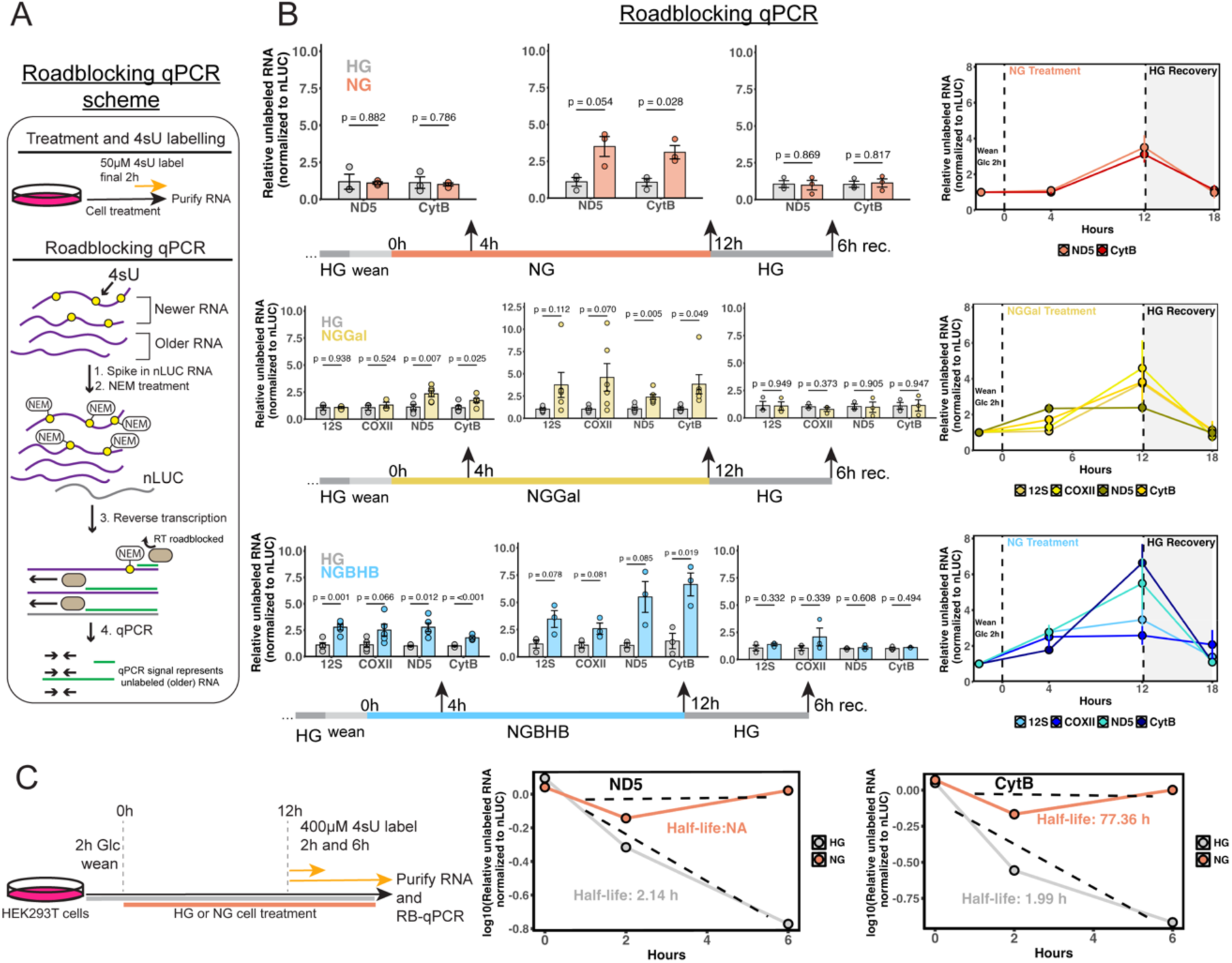
Mitochondrial RNA kinetics under different glucose deprivation conditions. **A)** General schematic of the roadblocking qPCR approach to measure RNA stability. **B)** Comparative roadblocking qPCR analysis of mitochondrial transcripts shows an increase in unlabeled RNA over time upon glucose deprivation. In NG- and NGBHB-treated cells subjected to a 6-h recovery in HG, unlabeled RNA levels returned to those observed in HG (n = 3-6 biological replicates, two-tailed Student’s *t*-test, data are means ± S.E.M. and associated *p* values). **C)** Representative mitochondrial RNA half-life measurements by roadblocking qPCR. Each dot on the plots represents the average of 3 biological replicates at different 4sU pulse times after a 12 h treatment (line-of-best-fit is shown as a dotted line and half-life values are displayed).

We observed a higher amount of unlabeled RNA upon glucose deprivation for mitochondrial transcripts ND5 and CytB after 12 h in all conditions (Fig. 2B). This implies a slowdown of mtRNA degradation and stabilization of older transcripts. Interestingly, while NG did not lead to accumulation of older RNA until 12 h into the treatment, NGGal and NGBHB treatment as short as 4 h showed a significant buildup of unlabeled RNA compared to HG (Fig. 2B). Additionally, we saw the highest degree of stabilization for ND5 and CytB transcripts in NGGal and NGBHB conditions compared to HG, while COXII was stabilized to a lesser degree and 12S ribosomal RNA only showed slight stabilization in the NGBHB condition compared to HG (Fig. 2B). To ask whether this decrease in mtRNA turnover was reversible, we cultured cells for 12 h in each condition before swapping the media back to high (25 mM) glucose without BHB for 6 h, followed by 4sU labelling during the final 2 h of growth. Strikingly, the unlabeled RNA at the end of the 6 h swap back to HG matched that of the HG control, indicating that mtRNA turnover is highly sensitive to the availability of glucose and recoverable on a relatively short time scale (Fig. 2B). In addition, we noted a similar accumulation of unlabeled RNA after a 12 h NG growth compared to HG in HepG2 cells, demonstrating a conserved mtRNA stabilization response to glucose deprivation (Fig. S2A).

The higher roadblocking qPCR signal in glucose-deprived cells (Fig. 2B) could be due to slower mitochondrial transcription (see the next section), i.e., less 4sU incorporation into mtRNA and thus less roadblocking during the RT step. To rule out changes in the rate of mitochondrial transcription as the cause of unlabeled RNA accumulation, we performed a complementary roadblocking qPCR experiment in which cells were starved of glucose for longer (12 h) prior to the 4sU addition, with 4sU-labeled samples analyzed at the 0, 2 and 6 h marks post-4sU addition and signal compared to the 0 h timepoint in each given condition (immediately before 4sU addition). This approach gives time for mitochondrial transcription rate to reach equilibrium under the given metabolic condition before the 4sU introduction, allowing the decay in signal to represent mitochondrial turnover alone. We noted the expected decrease in signal for the ND5 and CytB transcripts in HG, indicative of mtRNA decay (Fig. 2C and S2B). Linear regression of log-transformed relative values yielded half-life measurements of ~2 h for each transcript, in line with recent reports of mtRNA half-lives^12^. Strikingly, we did not observe the same decay pattern for either transcript in cells cultured in NG and extremely extended half-lives for each transcript (Fig. 2C and S2B). As a control, we measured the decay of the mitochondrial 12S ribosomal RNA in HG, which is expected to display a much longer half-life (~15 h) compared to the coding transcripts (Fig. S2C)^12,26^. These results establish that mitochondrial transcripts are stabilized during glucose deprivation and that roadblocking qPCR can accurately measure these changes.

### Mitochondrial transcription slows upon glucose deprivation

The surprising result that mtRNA turnover in cells slowed (older mtRNAs accumulated) but the steady-state pools of mtRNAs remained the same or slightly decreased upon glucose removal implies that mtRNA synthesis must slow down under these conditions. To confirm this experimentally, we employed pulse-chase immunofluorescence imaging with staining against the uridine analog bromouridine (BrU). BrU is incorporated into mtRNA during a short “pulse” of the analog and appears as puncta within the MRG upon anti-Br(d)U antibody staining (Fig. 3A). BrU in the media is then replaced with a large excess of uridine; thus, any *de novo* transcripts made during this “chase” phase will no longer contain BrU and be invisible in the anti-BrU channel. Two pieces of information can be extracted from such a pulse-chase assay: (i) anti-BrU signal intensity at the end of the pulse gives a sense of transcription rate (higher signal corresponds to higher rate of nucleotide addition); (ii) anti-BrU signal remaining after the chase informs on the composite rate of transcription initiation and RNA degradation (higher persisting anti-BrU signal in the presence of uridine competitor suggests slower transcription initiation and/or slower BrU-labeled RNA degradation). The difference between the two values yields an estimated “chase rate”, which represents the rate of RNA degradation.

**Figure 3.**
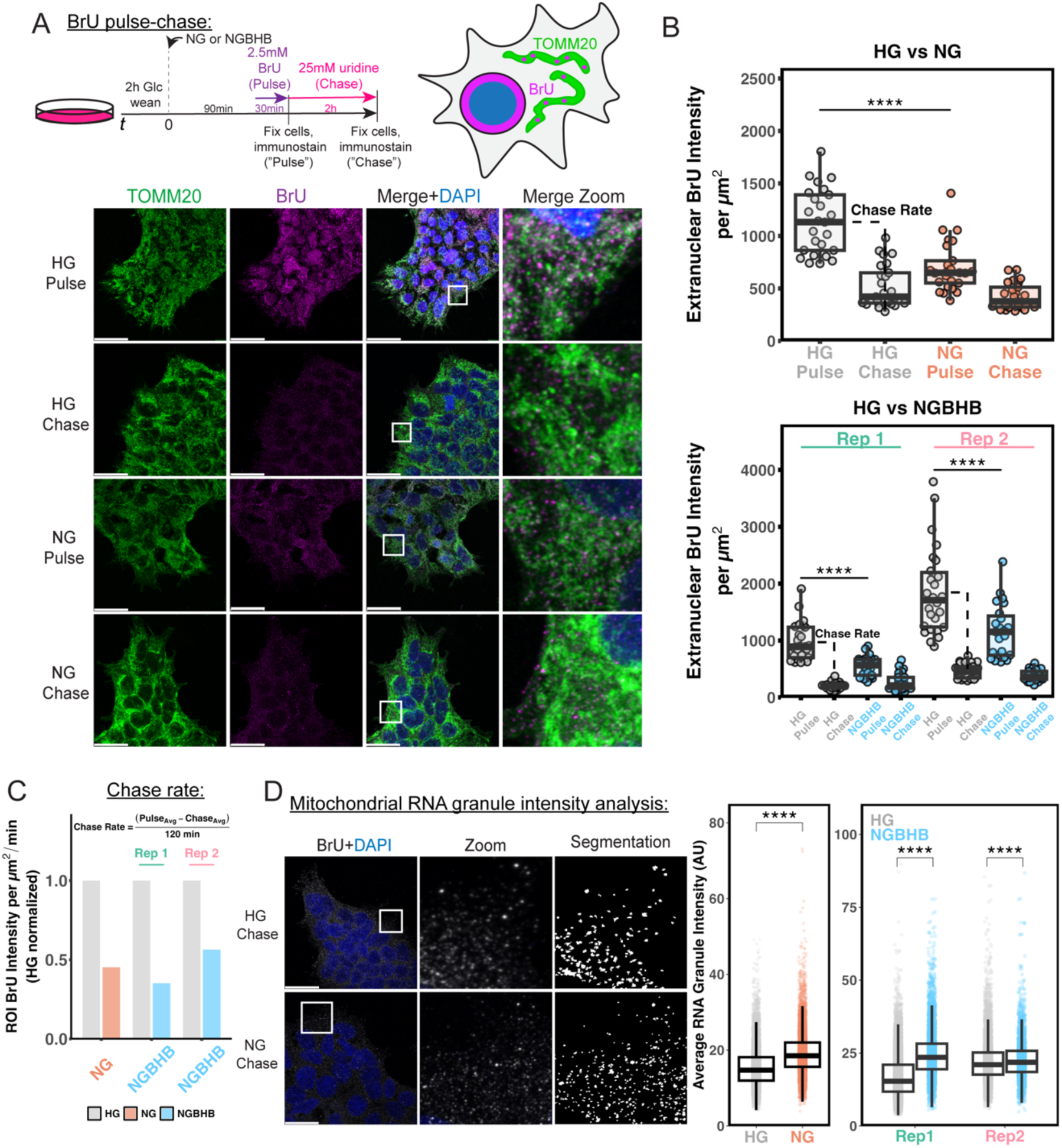
Pulse-chase experiments to visualize changes in mitochondrial transcription rate upon glucose deprivation. **A)** Schematic of a bromouridine (BrU) pulse-chase assay (left) and staining (right). HEK293T cells were cultured in HG, NG, or NGBHB for 90 min before a 30-min BrU pulse, after which portion of cells was fixed for immunofluorescence (“pulse” samples) to assess transcription rate. Remaining cells were then chased with 25 mM uridine for 2 h and fixed for immunofluorescence (“chase” samples) to assess RNA degradation rate. Representative images (bottom) are maximum intensity projections for HG vs. NG treated cells stained against TOMM20 (mitochondria) and BrU (newly synthesized RNA). Scale bar = 30 µm. **B)** Quantification of extranuclear BrU intensity per cell for pulse and chase samples of HG vs NG (top), and HG vs. NGBHB (bottom, \*\*\*\**p*<0.0001, one-way ANOVA with Tukey’s multiple comparisons correction; n=50 cells per replicate, with 25 cells in each condition quantified, data represents individual cells and bars represent median, boxes represent the 1^st^ and 3^rd^ quartiles, and whiskers are no further than 1.5 times the interquartile range). **C)** Quantification of BrU intensity chase rate, calculated as (average pulse – average chase)/120 min using data from panel B NG and NGBHB samples. **D)** Maximum intensity projection chase images for HG and NG treated cells, and segmented mitochondrial RNA granules (left, MRGs) for quantification of average pixel intensity per MRG from samples in panel B (right, \*\*\*\**p*<0.001, Wilcoxon rank sum test, data represent individual MRGs and box representation as above).

We chose to compare the HG to NG and NGBHB treatments due to the NGBHB’s pronounced mtRNA turnover phenotype on the shortest timescale (Fig. 2B). Briefly, the cells were cultured in HG or NGBHB for 90 min, then pulsed with BrU for 30 min, followed by a 2 h chase with uridine. Cells were immunostained and imaged using confocal microscopy both after the pulse and at the end of the chase (Fig. 3A). From Z-projections, we used image segmentation to quantify extranuclear BrU intensity per cell (Fig. S3A). We observed that NG and NGBHB-cultured cells incorporated significantly less BrU into their mtRNA during the pulse than HG-cultured cells, indicative of slower transcription in after 90 min of treatment (Fig. 3B, note: NGBHB replicates 1 and 2 were stained slightly differently, so data are shown separately for each replicate). Despite this difference in the starting BrU-labeled mtRNA, both HG- and NG/NGBHB-grown cells “chased” to a similar level (Fig. 3A, 3B and S3B). This suggests that mtRNA turnover (composite rate of *de novo* synthesis and degradation) slowed in NG and NGBHB, as indicated by the “chase rate” calculated as the difference between the average pulse and chase value in a given condition (Fig. 3C). From segmented regions of interest (ROIs), we observed no changes in translocase of outer mitochondrial membrane (TOMM20) signal across these conditions, validating that our anti-BrU signal derived from generally equal mitochondrial content (Fig. S3C).

These data complement the increased mtRNA stability in NG and NGBHB vs HG gleaned from 4sU roadblocking qPCR analysis (Fig. 2B) and establish that mitochondrial transcription rate is lower in an NG and NGBHB vs HG culture. As an additional support for slowed mitochondrial transcription, we saw a downward trend for the LSP1 primer synthesis by RT-qPCR at 12 h of NGBHB treatment (Fig. S3B). A recently characterized mitochondrial non-coding RNA, the 7S RNA, has been shown to directly inhibit POLRMT in cells and *in vitro*^27^. To rule out the possibility of mitochondrial transcription inhibition by the 7S RNA, we measured abundance of the mitochondrial 7S RNA with qPCR and found no difference in 7S RNA levels between HG and NGBHB conditions (even at 12 h of treatment), likely ruling the 7S RNA out as a cause for the observed decrease in transcription rate and further evidence of stabilized RNA pools (Fig. S3B).

When analyzing the chase images, we noticed that the NG and NGBHB images contained particularly bright MRGs compared to HG treatment, even though whole-cell anti-BrU intensity was “chased” to the same level overall in the two conditions. To capture this further, we segmented the chase images to measure the average pixel intensity per MRG (Fig. 3D, left and S3E). While many MRGs in HG and NGBHB chased to the same low intensity, there was a significant remaining bright population of MRGs in the NG and NGBHB chase samples (Fig. 3D, right). We hypothesize that these bright MRGs could be the indicators of a slower flux of newly synthesized RNAs out of MRGs or the recently described “storage” granules for mtRNAs during stress, such as pharmacological inhibition of mitochondrial transcription or toxic RNA buildup^28,29^. Taken together, these data support slowed mtRNA synthesis and potential changes to mtRNA flux from MRGs or mtRNA sequestration through metabolic stress (e.g., scarcity of glucose).

### Mitochondrial degradosome protein levels and localization to mitochondria are maintained during glucose deprivation

To determine if slowed mtRNA turnover in glucose-deprived cells was caused by altered levels of the proteins that dictate mtRNA stability and degradation, we first performed whole-cell western blot analysis of these proteins. Specifically, we blotted against the mtRNA chaperone LRPPRC and proteins important for mtRNA degradation (G-Rich RNA sequence binding factor 1, GRSF1; ATP-dependent RNA helicase SUPV3L1, SUV3; Polynucleotide Phosphorylase, PNPT1). None of these proteins showed any changes when subject to 12 h glucose deprivation compared to HG (Fig. S4A).

We next visualized the localization of the mitochondrial degradosome proteins SUV3, GRSF1, and PNPT1 in a 12 h NG treatment compared to HG. To accomplish this, we used a similar BrU labeling to mark mtRNAs as in the previous section, followed by immunostaining against BrU, TOMM20, and each degradosome component (Fig. 4A). We did not observe any obvious changes in protein localization, as shown in example line scans from each sample (Fig. 4B), with each degradosome component still localized to mitochondria and with mtRNA. Together, these findings indicate that the mtRNA stabilization phenotype is not caused by the changes in levels or localization of mitochondrial degradosome proteins. Rather, the mtRNA turnover appears to slow down through a universal energy-sensing mechanism.

**Figure 4.**
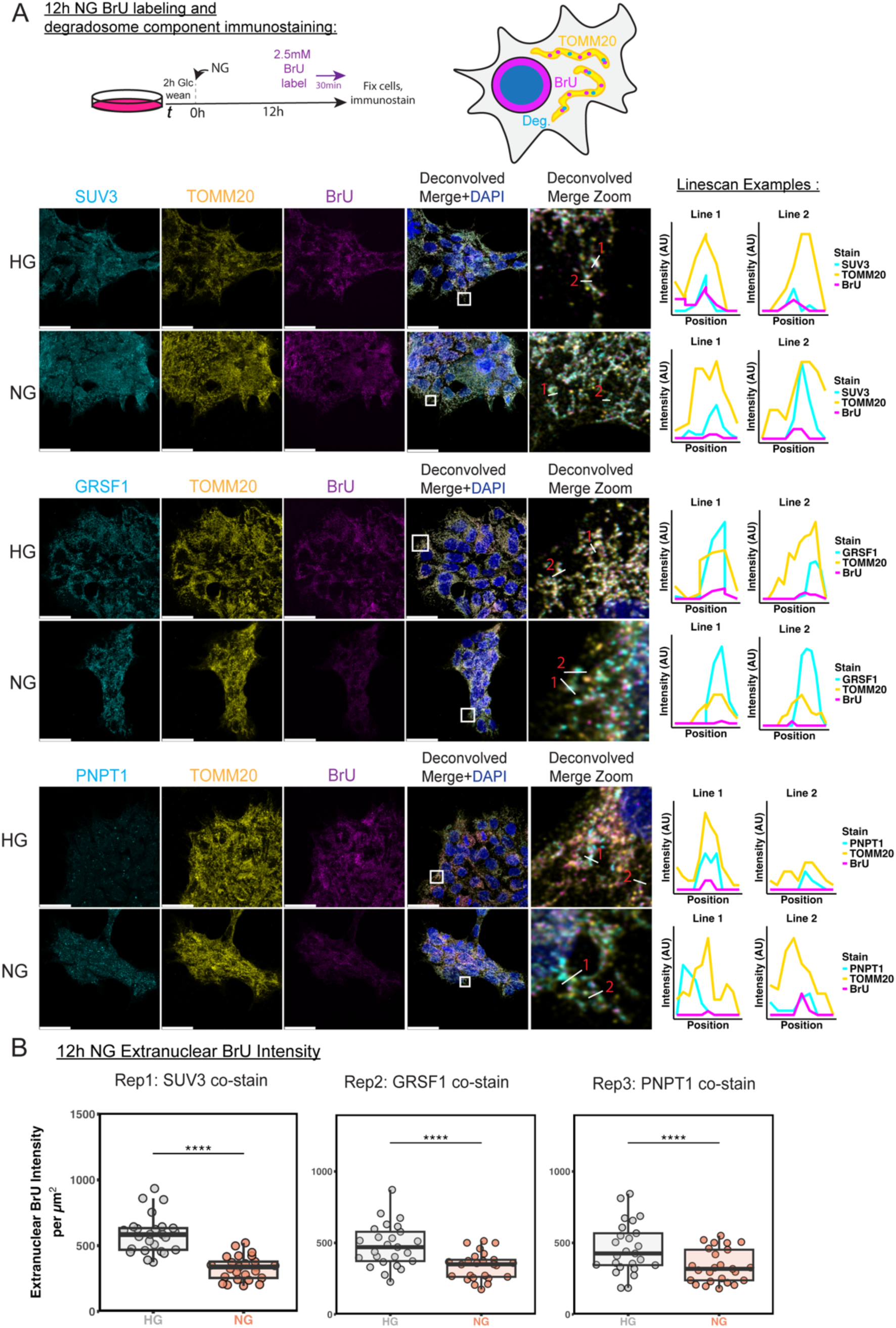
Mitochondrial degradosome proteins localize to mitochondria. **A)** Schematic of a bromouridine (BrU) incorporation assay (left) and staining (right) of TOMM20, BrU, and mitochondrial degradosome components (“Deg”). HEK293T cells were cultured in HG, or NG for 11.5 h before a 30-min BrU pulse, after which portion of cells was fixed for immunofluorescence. Representative images are shown as raw channel maximum intensity projections and a deconvolved merge maximum intensity projection for 12 h HG vs NG treated cells stained against TOMM20 (yellow, mitochondria), BrU (magenta, newly synthesized RNA), and mitochondrial degradosome components SUV3, GRSF1, and PNPT1 (Cyan). Scale bar = 30 µm. To the right of each image set are two example line-scans representing raw image intensities for each channel. **B)** Quantification of extranuclear BrU intensity per cell for 12 h HG vs NG BrU pulse samples in A (\*\*\*\**p*<0.0001, two-tailed Student’s *t*-test; n=50 cells per replicate, with 25 cells in each condition quantified; data represents individual cells and bars represent median, boxes represent the 1^st^ and 3^rd^ quartiles, and whiskers are no further than 1.5 times the interquartile range).

Using the same segmentation scheme for 90 min BrU-labeled MRGs as in the previous section, we measured the extranuclear BrU intensity per cell in these 12 h NG samples vs the respective HG control. We saw lower mitochondrial transcription rate in NG samples and unchanged TOMM20 intensities per ROI (Fig. 4B and S4B). These results along with those from Fig. 3B show a rapid and sustained drop but not a halt to mitochondrial transcription under glucose deprivation.

### Mitochondrial transcription rate mirrors mitochondrial ATP levels

Given the quick (within ~90 min) slowdown in mtRNA production upon glucose removal and the sustained (at 12 h) drop in transcription, we wondered if this was tied to mitochondrial ATP pools. Human mitochondrial transcription is hypothesized to be especially sensitive to local [ATP], and transcription initiation rate would reflect [ATP]-dependence because the first three nucleotides in mitochondrial transcripts are all adenosines. Indeed, *in vitro* studies with purified human mitochondrial transcription initiation proteins showed slower initiation of RNA synthesis at lower [ATP]^6^. Moreover, local [ATP] in the mitochondrial matrix is expected to directly affect the rate of mtRNA degradation because the helicase SUV3 that unwinds the double-stranded RNA (dsRNA) and RNA-DNA hybrids for exonuclease cleavage is ATP-dependent^13^.

To assess mitochondrial ATP content after glucose removal, we turned to a previously characterized, genetically encoded and mitochondrially targeted ratiometric ATP sensor^30^. The sensor contains a circularly permuted superfolder GFP (sfGFP) between two ATP binding helices of *Bacillus subtilis* F_o_-F_1_ ATPase, which upon binding ATP causes an increase in sfGFP fluorescence (Fig. 5A). The C-terminus of this sensor encodes a spectrally separate red fluorescent protein miRFP670nano, which acts as a control for sensor concentration and allows the ratio of green-to-red fluorescence intensity to report on mitochondrial ATP content (Fig. 5A).

**Figure 5.**
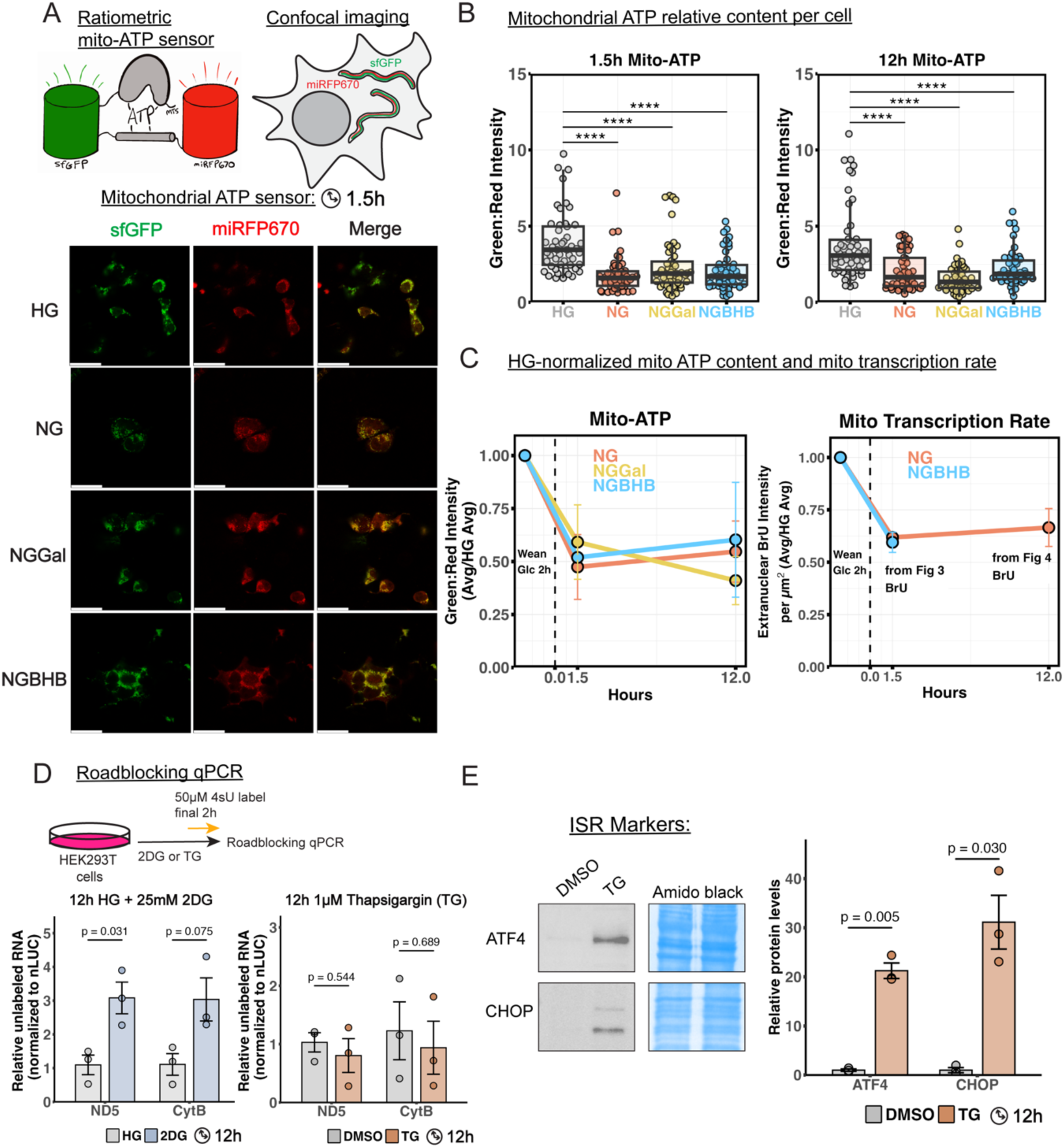
Mitochondrial ATP content reflects mitochondrial transcription rate. **A)** A cartoon showing the mitochondrial ATP sensor on the pAAV-CAG-iATPSnFR2A-95A-A119L-miRFP670nano3 plasmid (Addgene # 209723), which uses a circularly permuted sfGFP as a sensor between two ATP binding helices of *Bacillus subtilis* F_o_-F_1_ ATPase, and miRFP670 as a control (left). The ATP sensor is targeted to mitochondria with a mitochondrial targeting sequence (right). Representative 2D images of mitochondrial ATP sensor imaging in HEK293T cells during treatment for 1.5 h in glucose deprivation conditions (bottom). **B)** Quantification of mitochondrial ATP per cell as the ratio of green:red intensity for samples treated in each condition for 1.5 and 12 h (\*\*\*\**p*<0.0001, one-way ANOVA with Tukey’s multiple comparisons correction; n=50 cells per condition, with 25 cells in each replicated from 2 biological replicates; quantified points are individual cells and bars represent median, boxes represent the 1^st^ and 3^rd^ quartiles, and whiskers are no further than 1.5 times the interquartile range). **C)** Summarized timeline results from mitochondrial ATP measurements (left) and BrU pulse assays (right) as each conditions average value normalized to the respective HG value. Data represents the mean and S.E.M. for each sample. **D)** Experimental schematic and roadblocking qPCR analysis of ND5 and CytB in HEK293T cells treated in 10 mM 2-deoxyglucose (2DG) for 12 h in the presence of high glucose or 1 µM thapsigargin (TG) for 12 h, showing RNA stabilization with 2DG treatment (n=3 biological replicates, two-tailed Student’s *t*-test, data is mean ± S.E.M. with associated *p* values). **E)** Representative western blots of ATF4 and CHOP protein levels in HEK293T cells treated with 1 µM thapsigargin (TG) for 12 h compared to DMSO treated (left). Quantification of blots is shown on the right (n=3 biological replicates, normalized to the average HG or DMSO values respectively, two-tailed Student’s *t*-test with associated *p* values).

Using confocal live-cell 2D imaging and segmentation of each channel, we quantified the mitochondrial [ATP] per cell for our glucose deprivation conditions compared to HG at the 2 h and 12 h treatment marks (Fig. 5A and S5A). We saw a significant drop in mitochondrial [ATP] in all conditions (Fig. 5B and S5B) at both timepoints with no significant differences between the glucose deprivation conditions. This was a surprising result because we expected BHB to boost mitochondrial [ATP], being a mitochondria-specific fuel. The sustained drop in mitochondrial [ATP] is an important finding, given that all glucose deprivation conditions lead cells to solely use OXPHOS for energy production (Fig. 1C), whereas HG-grown cells are both highly glycolytic and oxidative (Fig. 1C). It is possible that cells reliant on OXPHOS upregulate export of ATP from mitochondria, especially in conditions where glycolysis is shut off. Strikingly, when we compared a normalized transcription rate and mitochondrial [ATP] over our two timepoints, we saw that both dropped to a similar relative level (~50% mitochondrial [ATP] and ~60% mitochondrial transcription compared to HG) and persisted there, providing further evidence that mitochondrial transcription can respond to changes in metabolism and potentially sense compartmental [ATP] (Fig. 5C).

Given the drop in mitochondrial [ATP], we next checked on the activity of the RNA helicase SUV3 inferred from immunostaining with the dsRNA specific antibody J2 of 12 h NG treated cells. We found a slight increase in extranuclear J2 intensity compared to HG, but not an increase on par with the degree of stabilization of mtRNAs in this condition (Fig. S5C). This could be due to the slowed transcription reducing the burden on SUV3 to unwind mitochondrial dsRNA. Most importantly, we did not see large dsRNA foci induced by long-term SUV3 ablation^31^, arguing that stabilized mtRNAs are likely still traversing through the mitochondrial central dogma in our glucose deprived conditions, albeit more slowly.

### Glycolytic insults but not ER-stress induced ISR signaling stabilizes mtRNA

Considering that glucose deprivation induces ISR signaling, we applied roadblocking qPCR to determine if glycolytic inhibition or direct induction of ISR signaling in nutrient abundance could recapitulate the mtRNA stabilization phenotype. To directly target glycolysis, we treated HEK293T cells grown in 25 mM glucose with 25 mM 2- deoxyglucose (2DG), which upon conversion to 2-deoxyglucose-6-phosphate can inhibit the glycolytic enzymes hexokinase and phosphoglucose isomerase^32,33^. We indeed found that a 12 h 2DG treatment caused the stabilization of ND5 and CytB transcripts seen in glucose deprivation conditions, while steady-state qPCR shows maintained overall transcript levels (Fig. 5C and S5C). These results support the notion that inhibition of glycolysis and potentially the resulting drop in mitochondrial [ATP] are closely tied to the mtRNA stabilization and transcription slowdown.

To test whether signaling upstream (e.g., ISR activation) of mtRNA synthesis or the downstream fate of mtRNA (translation) contributed to the observed mtRNA stabilization, we turned to pharmacological treatments that target each of these processes. To illicit direct ISR activation, we treated cells with 1 µM thapsigargin (TG), an inhibitor of the sarcoplasmic/ER Ca^2+^-ATPase (SERCA) required for pumping Ca^2+^ into the endoplasmic or sarcoplasmic reticulum^34^. TG treatment depletes ER Ca^2+^, leading to ER stress and the ISR^35^. Cells treated with TG did not show the ND5 or CytB transcript stabilization compared to a DMSO (vehicle)-treated control, despite activation of the UPR evident from the increased levels of ATF4 and CHOP, classic markers of ISR^21,36^ (Fig. 5C, 5D and S5D). Thus, we propose that the ISR coincides with but does not necessarily lead to mtRNA stabilization and mitochondrial transcription slowdown. To directly inhibit mitochondrial translation, we treated cells with chloramphenicol, an antibiotic specific for mitoribosomes^31^. We found that chloramphenicol treatment generally left mtRNA transcript lifetimes unaffected except for destabilization of ND5 (Fig. S5E).

### Changes in mitochondrial RNA synthesis and stabilization are generally independent of mtDNA replication

With the knowledge of mitochondrial transcription being absolutely required for mtDNA replication, we measured the relative mtDNA copy number (CN) under each glucose deprivation condition using qPCR. In agreement with past studies^23^, NG and NGGal did not change the mtDNA CN, but the NGBHB condition caused a ~2-fold increase in CN at 12 h compared to HG in HEK293T cells (Fig. 6A). We found that the NGBHB-induced CN increase persisted at 24 h in HEK293T cells but this effect was not observed in other tested cell types (HepG2, U2OS, and Mouse Embryonic Fibroblasts, MEF) (Fig. 6A). Furthermore, when HEK293T cells were treated with 6 mM BHB in the presence of high glucose (HGBHB), the increase in mtDNA CN was no longer observed, implicating *both* BHB and lack of glucose in the CN increase (Fig. 6A). Given that BHB is derived from fatty acids, we probed HEK293T cell mtDNA CN using a 12 h treatment without glucose and supplementation with one of the two most abundant fatty acids in humans, palmitic acid and oleic acid, and found no change in CN compared to HG (Fig. S6A). In light of the characterization of HEK293 lineage as a combination of liver, neuronal, and adrenergic cells^37^, and the literature implicating BHB as a brain fuel during fasting, we next treated SH-SY5Y, a typical cell line used as a neuronal model, and a differentiated neural spheroid model with a physiologically feasible concentrations of low glucose (1 mM)^38^ and 6 mM BHB for 12 h and observed no change in mtDNA CN compared to the HG control (Fig. S6B), further demonstrating that the NGBHB-induced increase in mtDNA CN is specific to HEK293T cells.

**Figure 6.**
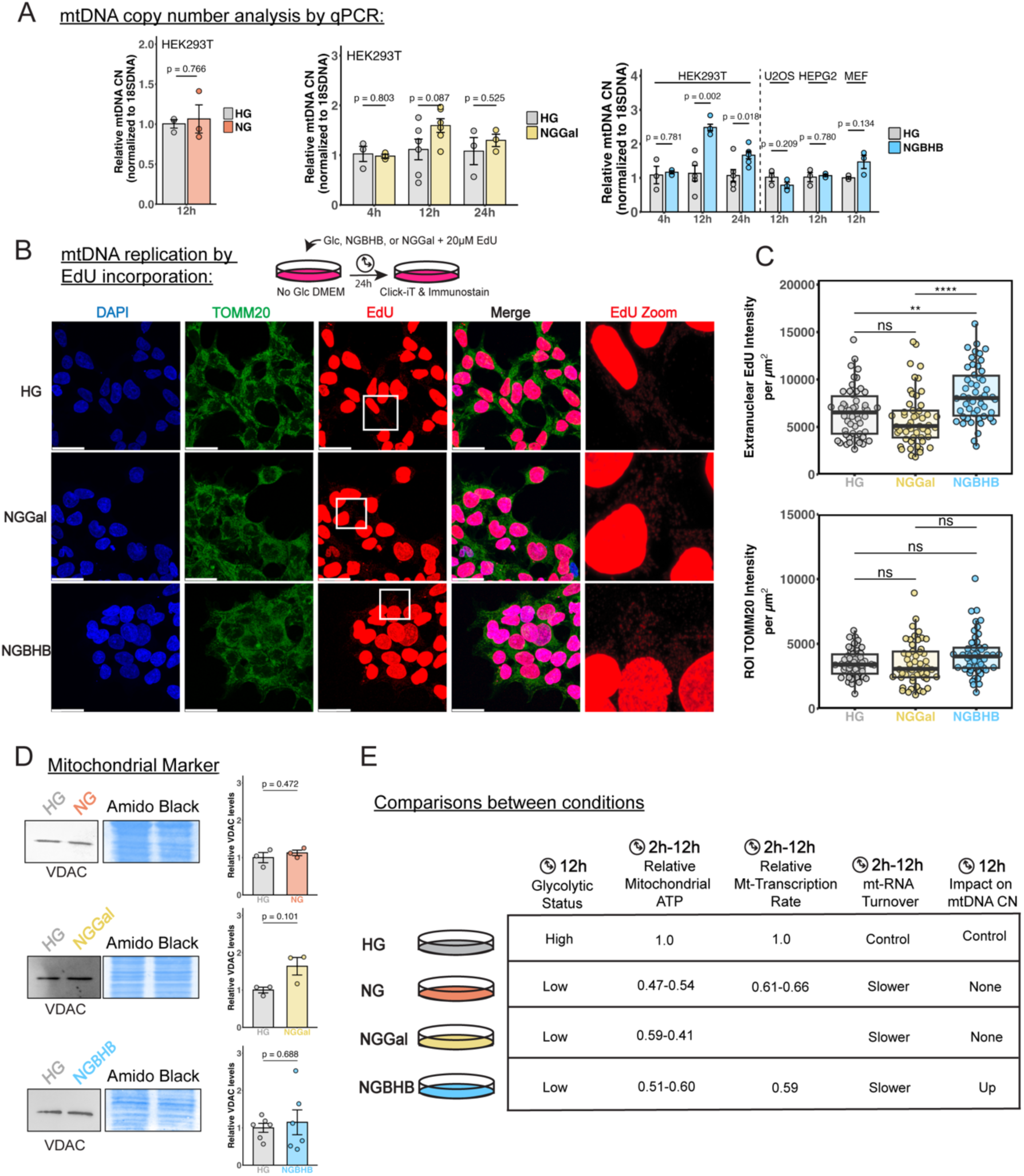
Combination of glucose deprivation and BHB supplementation increases mitochondrial DNA content in HEK293T cells. **A)** Quantitative PCR (qPCR) analysis of mtDNA copy number (CN) in high glucose (HG) and glucose deprivation conditions. mtDNA was detected with primers for CytB normalized to 18S DNA (n=3-6 biological replicates, two-tailed Student’s *t*-test, data is mean ± S.E.M. with associated *p* values). **B)** Schematic for EdU incorporation into newly synthesized DNA and immunostaining in HEK293T cells (top) and representative images as max projections (bottom; scale bar = 30 µM). **C)** Quantified extranuclear EdU intensity per cell for each condition and total TOMM20 signal per ROI as a control for mitochondrial content (right, ns, nonsignificant; \*\*\*\**p*<0.0001, \*\**p*<0.01, one-way ANOVA with Tukey’s multiple comparisons correction, n=50 cells: 25 cells each condition were quantified for 2 biological replicates, data represent individual cells and bars represent median, boxes represent the 1^st^ and 3^rd^ quartiles, and whiskers are no further than 1.5 times the interquartile range). **D)** Representative western blots of VDAC in HEK293T cells treated with glucose deprivation conditions 12 h compared to HG (left). Quantified blots are shown on the right (n=3 biological replicates, normalized to the respective average HG values, two-tailed Student’s *t*-test with associated *p* values). **F)** Summary of the findings for glucose deprivation’s effects on metabolism, mitochondrial [ATP], mitochondrial transcription rate, RNA stability, and mitochondrial DNA copy number.

To provide visual support for the increased mtDNA CN and thus increased mtDNA replication, we used metabolic labelling and immunofluorescence detection with a Click-iT 5-ethynyl 2’-deoxyuridine (EdU) Labelling kit to detect newly synthesized mtDNA (Fig. 6B). Briefly, HEK293T cells were cultured in the presence of 20 µM EdU (a synthetic nucleoside that is incorporated into newly synthesized DNA) for 24 h in the HG, NGBHB, or NGGal conditions followed by indirect immunolabelling, confocal imaging, image segmentation and quantification of extranuclear EdU signal per cell. Image analysis revealed a significant increase in newly synthesized mtDNA in the NGBHB condition compared to both the HG and NGGal (Fig. 6C and Fig. S6D), validating mtDNA CN measurements by qPCR (Fig. 6A). We found no significant difference in TOMM20 signal within our quantified ROIs, validating EdU intensity results (Fig. 6C). Importantly, western blot analysis of the voltage-dependent anion channel (VDAC) as a proxy of mitochondrial mass showed no differences after treatment with each glucose deprivation condition compared to HG (Fig. 6D), demonstrating that the increase in mtDNA CN in the NGBHB condition is independent of mitochondrial mass.

A recent study on mitochondrial nucleoid proteins found that inhibition of mitochondrial translation with chloramphenicol can stimulate mtDNA replication^39^, a response with a currently unknown mechanism but is hypothesized to be compensatory. Given the fact that the HGBHB condition did not show this response, we wondered if BHB imposes an added stress on the mitochondrial central dogma to elicit an increase in mtDNA CN. Recent reports demonstrated that BHB can post-translationally modify lysine residues on proteins in human cells as a non-canonical acylation and that modification is widespread. One study reports a primary target of this modification being the glycolytic enzyme aldolase (~39 kDa), with BHB modification inhibiting aldolase activity^40^, while others report that a majority of KBHB is found on nuclear proteins, with BHB modifying histones directly^41,42^. In support of this recent work, we saw widespread lysine-β-hydroxybutyrylation in HEK293T cell lysates treated with NGBHB for 12 h compared to HG (Fig. S6E). Additionally, using cell fractionation and histone extraction, we detected KBHB on histone-enriched samples using anti-KBHB western blots (Fig. S6F).

We analyzed RNA-seq data to identify differentially expressed genes between 12 h NGBHB and NG treated HEK293T cells. We found very few genes to be differentially expressed but an increase in expression for genes related to steroid and cholesterol synthesis (Fig. S6G and S6H), some of which can reside in mitochondria or the ER, while others are related to transcription and the peroxisome^43–49^. This trend was also recently reported in transcriptomics of mice livers from mice subject to a ketogenic diet^40^. Intriguingly, steroid and cholesterol metabolism involves both the ER and mitochondria, and a prior study has shown that mtDNA replication is “licensed” at mitochondria-ER contact sites^50^, suggesting a potential connection between steroid and cholesterol metabolism and mtDNA CN. The specificity of the mtDNA CN increase to HEK293T cells may potentially be attributed to HEK293 adrenal lineage^37^.

## Discussion

Glucose deprivation sets off a complex series of adaptations to keep cells alive, including breakdown of glutamine, a shift to an oxidative metabolism, sequestration of cytosolic RNAs, and inhibition of cytosolic translation except for a select set of stress-related messenger RNAs^16,51–53^. Here, we add to this complex picture by showing that mitochondria respond by reducing transcription of their genome and stabilizing mtRNAs to maintain steady pools of mtRNA, a phenotype that is quickly reversed with re-introduction of glucose (Figs. 1A, 2B-C, and 3B-C).

The reduction in mitochondrial transcription rate in response to glucose deprivation is rapid, as we noted a drop in mtRNA synthesis to ~60% compared to HG after 90 min, with the slower mtRNA synthesis sustained at 12 h (Figs. 3B, 4B, and 6F). Importantly, we found that mitochondrial [ATP] was lower in glucose deprivation conditions compared to HG and closely mirrored mitochondrial transcription rate across this timeline. This, combined with reliance on OXPHOS (Fig. 1C and 6F), points to an upregulated mitochondrial ATP export when glycolysis does not produce ATP and argues that mitochondrial transcription rate can be predicted based on the energetic status of the cell. The recovery in mtRNA turnover upon glucose reintroduction points towards a synchronized adjustment in the rates of mtRNA synthesis and degradation. We noted brighter MRGs under glucose deprivation and a slower “chase” of BrU-labeled mtRNAs (Fig. 3A and 3B); however, mtRNAs were still being processed (Fig. S2D), suggesting a potential slowdown in mtRNA movement out of MRGs, indicated by the accumulation of BrU-tagged RNAs in these granules. Additionally, the components of the mitochondrial degradosome also still localized to mitochondria (Fig. S4B) and preserved their activity (e.g., SUV3 in Fig. S5C), suggesting that all stages of the mtRNA lifecycle are sustained under glucose deprivation but slowed. This complements previous work showing that removal of glucose and replacement with galactose does not alter mitochondrial translation in the long term (48-72 h)^16,23^.

We propose a flux-based model for regulation of mtRNA kinetics in response to nutrient availability (Fig. 7). During nutrient abundance, mtRNAs are overproduced (compared to the nuclear-encoded OXPHOS mRNAs) and passed to a system of mitochondrial bodies or proteins (MRGs, LRPPRC, ribosomes, and the degradosome) with finite pools of proteins available. We propose that these proteins are limiting, leaving excess mature RNAs to be degraded (Fig. 7A). Under nutrient deprivation, ATP-dependent mtRNA synthesis slows down, producing fewer “excess” mtRNAs and matching more closely the finite protein capacity of the mtRNA bodies and chaperones. Excessive mtRNA transcription could thus be an adaptive process that acts as a buffer to cellular insults while metabolism adjusts to allow the cell to survive those insults.

**Figure 7.**
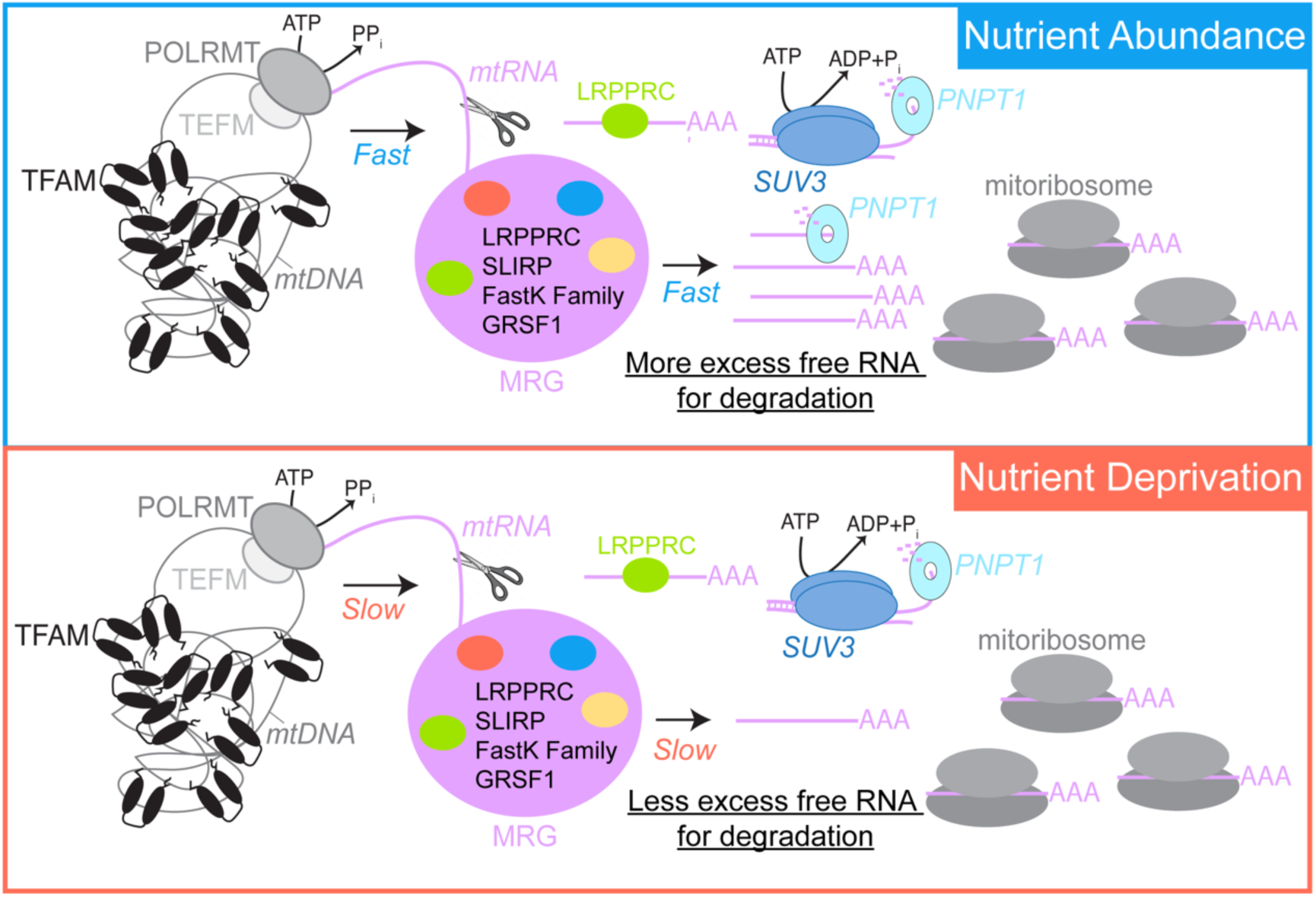
A flux-based model of mitochondrial gene expression changes under glucose deprivation. Overproduction of mitochondrial RNAs and movement through the mitochondrial RNA granule (MRG) in nutrient abundance leads to faster degradation rates (top), while slowing these processes during nutrient deprivation leads to fewer excess mtRNA produced and longer mtRNA lifetimes.

Our data differ from two recent studies showing new “bodies” formed by mtRNA under stress: one showing mitochondrial mRNA sequestered during long-term mitochondrial transcription inhibition with ethidium bromide or IMTB1^54^, and another demonstrating that long-term SUV3 ablation leads to formation of distinct foci containing dsRNA and mitochondrial rRNA^31^. We believe the lack of these phenotypes in our experiments is due to the harshness of the insult in each study, in which specific parts of the mtRNA life cycle were inhibited during nutrient excess, therefore not showing a concerted shift across all parts of the mtRNA lifecycle. In contrast, our data demonstrate that an intact mitochondrial central dogma can quickly adapt to an insult like nutrient deprivation without forming distinct mtRNA stress bodies. This work compliments a recent study in which ISR activation and subsequent cytosolic translation inhibition were shown to selectively allow local continued translation of mitochondrial proteins^55^. Our model predicts that the mitochondrial transcription system would be adapting to a slowed state to match this response.

## Materials and Methods

### Cell culture and transfection

HEK293T (ATCC CRL-3216), HepG2 (ATCC HB-8065), U2OS (ATCC HTB-96), and MEF (ATCC SCRC-1008) cells were maintained under 5% CO_2_ at 37 °C in DMEM medium containing 25 mM glucose and 2 mM glutamine (11965092, Gibco) supplemented with 10% FBS (A5256701, Gibco) and 1% penicillin/streptomycin (10378016, Gibco). SH-SY5Y (ATCC CRL-2266) cells were maintained in DMEM/F12 media (11320033, Gibco) supplemented with 10% FBS, 1% penicillin/streptomycin, and 1X Glutamax (35050061, Gibco). For cell passaging, cells were first washed in warmed PBS (10-010-023, Gibco) and then incubated in warmed ATV solution or (25200056, Gibco) for 5-10 minutes. The cells were collected, added to complete warmed DMEM to inactivate the ATV solution, and pelleted with a spin at 400-1500 x g for 3-8 minutes. The medium was aspirated, and the cells were resuspended in fresh complete DMEM for cell counting and plating. Cells were maintained for a maximum of 20 passages. Cells were checked for mycoplasma contamination using a Mycoplasma PCR Detection Kit (ab289834, Abcam).

The motor neurons were differentiated as described by Kawada et al. Undifferentiated human iPSCs were maintained in culture in iPSC culture medium (mTeSR, STEMCELL) until ~70% confluency in a 6-well plate, followed by PBS wash and dissociation with TrypLE Select (Thermo Fisher Scientific). Cells were resuspended in iPSC culture medium with 10 μM Y-27632 dihydrochloride (Tocris) and seeded at 40,000 cells/well in low-attachment 96-well U-bottom plates (Corning) medium containing 10 μM Y-27632 dihydrochloride. Differentiation was initiated using DMEM/F12 (Gibco) supplemented with 15% Knockout Serum Replacement (Thermo Fisher Scientific), GlutaMAX (Thermo Fisher Scientific), non-essential amino acids (Thermo Fisher Scientific), 10 μM SB-431542 (Sigma-Aldrich), and 100 nM LDN-193189 (Sigma-Aldrich) for 2 days, followed by a gradual transition to N2 medium over 8 days with 1 μM retinoic acid (Sigma-Aldrich), 1 μM Smoothened Agonist (Sigma-Aldrich), 5 μM SU5402 (Sigma-Aldrich), and 5 μM DAPT (Sigma-Aldrich). On day 12, spheroids were transferred to a tissue culture plate in maturation medium, 20 ng/mL BDNF (Preprotech). Prior to transfer, spheroids were embedded in Matrigel (1:40 in DMEM/F12) in a 6-well plate using wide-bore pipette tips. Medium was changed every 2–3 days, and axon networks formed within 1 week.

### Glucose starvation experiments, drug treatments, and cell counting

For starvation experiments, no-glucose DMEM containing 2 mM glutamine (11966025, Gibco) was supplemented with 1% penicillin/streptomycin and 10% dialyzed FBS. FBS was dialyzed overnight at 4 °C against prepared PBS (137 mM NaCl, 2.7 mM KCl, 100 mM Na_2_HPO_4_, 18 mM KH_2_PO_4_, pH 7.4, filter sterilized with 0.22 µM filter). Glucose (cat. #D16-500, Thermo Fisher Scientific), galactose (cat. #A12813.30, Thermo Fisher Scientific), β-hydroxybutyrate (BHB, cat. #298360, Sigma Aldrich), and 2-deoxyglucose for supplementation were dissolved in PBS and filter sterilized using a 0.22 µM filter. Complete no-glucose DMEM (Gibco, cat. #11966025) was used to make different treatment solutions on the day of each experiment. To prepare high-glucose treatment medium the no-glucose medium was spiked to a final concentration of 25 mM glucose. No glucose, BHB, and galactose treated cells were weaned off glucose for 2 hours at 5 mM glucose before complete removal of glucose from culture media with or without alternative carbon source (6 mM BHB or 5 mM galactose) for the specified times. For live-cell imaging, cells were cultured in DMEM without glucose, glutamate, or pyruvate, and phenol red (Gibco, cat. #A1443001) supplemented with FBS dialyzed against PBS, and spiked with 1X GlutaMAX (Gibco, cat. #35050-061) supplement and spiked with the treatments above. Cells were treated at ~80% confluence by aspiration, washing with warm PBS, and then adding the warm treatment media.

Cell counting experiments were performed by seeding ~1500 cells per well in 80 µl of complete DMEM in a 96-well dish. Cells were allowed to adhere to the surface of the dish for ~8 h before aspirating the cells, washing them in the respective treatment media, and then incubating in 100 µl of the treatment media. Every 12 h cells to be counted were aspirated, incubated in Trypsin-EDTA (0.25%) for 15 min, stained with trypan blue (Gibco, cat. #15250-061), and counted using a Countess 3 Automated Cell Counter (ThermoFisher Scientific). Total cells and the percent of dead cells as reported by the trypan blue staining were recorded. For longer treatments, medium was replaced every 12 h.

### RNA-seq analysis

Cells were seeded and grown in 6-well plates until 70-80% confluency, at which point cells were subjected to starvation experiments as previously described. Cells were then aspirated, and 500 µl of Trizol Reagent (Invitrogen) was added to each well. The cells were then mixed on a rocking plate at 4 °C for 10 min and collected in centrifuge tubes. To each tube, 200 µl of chloroform was added, the samples were vortexed, and then spun at 18,407 x g for 12 min at 4 °C. The supernatant of each was collected, and RNA was purified with the Zymo RNA Clean and Concentrator Kit-5 (cat. #R1013), including the kit’s provided DNase treatment. RNA was eluted in water, and its concentration was determined by 260 nm absorbance or the Quant-iT™ RNA Assay Kit (cat. #Q33140, Thermo Fisher Scientific). RNA samples were submitted to the UCSD IGM Genomics Center for sequencing preparation, where total RNA was assessed for quality with an Agilent Tapestation 4200, and samples with an RNA Integrity Number (RIN) greater than 8.0 were used to generate RNA sequencing libraries. The Illumina® Stranded mRNA Prep (Illumina, San Diego, CA) was used to generate libraries from 500 ng of RNA, following manufacturer’s instructions. Resulting libraries were multiplexed and sequenced with 150 basepair (bp) Paired End reads (PE150) to a depth of approximately 25 million reads per sample on an Illumina NovaSeq X Plus. Samples were demultiplexed using bcl2fastq Conversion Software (Illumina, San Diego, CA).

Raw RNA-seq read quality was tested with FastQC (v0.12.0) and reads were trimmed with Trimmomatic (v0.39)^56^ using the following settings: LEADING:3 TRAILING:3 SLIDINGWINDOW:4:15 MINLEN:36. Paired reads were then aligned to the human reference genome (GRCh38.p14) primary assembly from GENCODE using STAR (v2.7.11b)^57^. The raw counts table was then prefiltered genes containing less than 10 total reads were removed before differential gene expression analysis was carried out in R using the DEseq2 package^58^ with default settings. For volcano plots fold changes were log_2_ transformed and plotted on the x-axis, while corrected p-values were −log_10_ transformed and plotting on the y-axis. Differentially expressed genes were determined as genes with an adjusted *p*-value <0.05 and a fold-change of 1.3 or greater. Gene ontology (GO) enrichment analysis of differentially expressed genes was carried out using the enrichR package^59^. Heatmaps were created using the ComplexHeatmap package. Lists for nuclear and mitochondrial encoded OXPHOS subunits were obtained from the Mitocarta 3.0^60^. FPKM analysis was carried out from the raw counts table.

### Oxygen Consumption and ECAR Measurements

Basal OCR and ECAR measurements were performed on a Seahorse XFe96 Analyzer (Agilent Technologies) using the Seahorse CF Cell Mito Stress Kit (Agilent Technologies). The Seahorse microplate wells were coated with poly-D-lysine (PDL) and 20,000 cells were seeded into each well and allowed to adhere for ~8 h. The wells were then washed once with treatment media listed above and finally treated in 100 µl of respective treatment media for 12 h at 37 °C and 5% CO_2_. The cells were then aspirated, washed, and swapped into the prepared Seahorse media (Agilent Seahorse media supplemented with 2 mM glutamine and either 25 mM glucose, 6 mM BHB, or 5 mM galactose). The cells were then incubated at 37 °C in a non-CO_2_ incubator for 1 hour before basal OCR and ECAR measurement. Final inhibitor concentrations used in the assay were 1 µM oligomycin, 1 µM carbonyl cyanide-4-phenylhydrazone (FCCP), 0.5 µM antimycin A, and 0.5 µM rotenone. A run report was generated using Wave Software (Agilent Technologies). Basal OCR for each well was determined as the average of the first three OCR measurements with subtraction of the average non-mitochondrial respiration. Basal ECAR was calculated as the average of the first three ECAR measurements.

### RT-qPCR, roadblocking qPCR, and relative mtDNA copy number determination

RNA was purified as previously mentioned for RNA sequencing. Complementary DNA (cDNA) synthesis was performed with 1 µg of purified RNA, Superscript II Reverse Transcriptase (Invitrogen, cat. #18064014) and random hexamer primers (Thermo Scientific, cat. #SO142) using standard protocols. The final 20 µl cDNA sample was diluted 1:10 with ultrapure water for use in qPCR. qPCR experiments were carried out with the PowerUP SYBR Green Master Mix (Thermo Fisher Scientific), using 1 µl of cDNA per reaction in 10 µl total volume with primers listed in Table 2. Experiments were completed using a QuantStudio3 Real-time PCR System with three biological replicates and either three or four technical replicates. Relative RNA levels were calculated with the ΔΔC_T_ method using TATA-binding protein (TBP) as an internal control (Primers are listed in Supplementary Table 4). Data were analyzed per plate, using 4 technical replicates per gene. Values were only excluded if determined to be an outlier using the Grubbs’ Test with a significance level of 0.05 (Graphpad).

Nanoluciferase (nLUC) RNA was synthesized using the HiScribe T7 High Yield RNA Synthesis Kit (New England Biolabs) with Addgene #199755 as a transcription template (see Supplemental Table 4 for primers). RNA was purified with an RNA Clean and Concentrator-5 kit (Zymo). nLUC RNA was then poly-A tailed using *E. coli* Poly(A) Polymerase (New England Biolabs) and repurified using the Zymo RNA Clean and Concentrator-5 kit. Cell treatments and RNA purifications were carried out as mentioned previously in the “Glucose starvation experiments” section above, except that cells were treated with 50 µM 4sU (Sigma Aldrich) for 2 h before RNA extraction with Trizol Reagent (ThermoFisher Scientific, cat. #15596026). For RNA half-life measurement samples, 400 µM 4sU was added to cells for 2 or 6 h, while the control 0 h timepoint was not treated with 4sU before purification of RNA as above. For half-life measurements, 500 ng of purified RNA was used for subsequent steps. For 4sU-labeled RNA alkylation, we followed the procedure by Watson and Thoreen^61^. Specifically, 1 µg (condition comparisons) or 500 ng (half-life measurements) of purified RNA was first denatured at 65 °C for 5 min in a total of 11 µl water, before addition of 10 µl of reaction buffer (final concentration 50 mM Tris-HCl pH 8.0, 1 mM EDTA), 1 µl of 0.2 ng/µl nLUC RNA, and 3 µl of 50 mg/ml NEM (prepared fresh in 100% ethanol). The alkylation reaction was incubated at 42 °C for 90 min before RNA purification with the Zymo RNA Clean and Concentrator-5 kit and elution with 20 µl of water. 6 µl of eluted RNA was then used for cDNA synthesis by Superscript II Reverse Transcriptase with PolydT_15_ and 12S RNA primers. qPCR and analysis were carried out as above. For RNA half-life measurements, the 2 h and 6 h samples were compared to their respective 0 h timepoint using the ΔΔC_T_ method. The ΔΔC_T_ method fold-change values were log_10_-transformed and linear regression was performed in R using the “lm” function. Half-lives were estimated using the provided slope and taken as the time to reach half of the 0 h timepoint’s ΔΔC_T_ fold-change value, (log10(0.5*0h)-log10(0h)/slope.

For mtDNA copy number analysis, cells were treated in 6-well dishes before harvest in ice-cold PBS. Total DNA was then extracted using a PureLink™ Genomic DNA Mini Kit (Thermo Fisher Scientific). qPCR reactions were carried out as above using 4-10 ng of total DNA template per reaction. Relative mtDNA copy number was calculated with the ΔΔC_T_ method using CytB primers for mtDNA detection and primers for 18S DNA as an internal control.

### BrU pulse-chase, EdU incorporation, and immunostaining

HEK293T cells were seeded on poly-D-lysine coated coverslips (Gibco) and grown to ~75% confluence and treated with HG, NG, or NGBHB as described in the “Glucose starvation experiments” section above. BrU and uridine stocks (250 mM and 1.25 M) were prepared in PBS, filtered, and stored at −80 °C. For BrU pulse-chase experiments, we adapted the protocol from Jourdain et al.^62^ At 90 min of starvation, the medium was supplemented with 5 mM BrU for 30 min. The medium was then aspirated, and cells were washed twice in 37 °C PBS. Chase samples were then incubated with either HG, NG, or NGBHB media containing 250 mM uridine for 2 h while pulse samples were fixed in a solution of 4% formaldehyde in pre-warmed PBS for 15 min at room temperature (RT). After the 2 h chase, samples were aspirated, washed and fixed as pulse cells above. NGBHB replicate 1 and its associated HG sample (Fig. 3B) were stained with wheat germ agglutinin-640R (WGA, Biotium) at this stage, which we did not image as part of confocal microscopy, whereas NGBHB replicate 2 and the NG samples were not stained with WGA. For this reason, we plotted and analyzed NGBHB replicates 1 and 2 separately (Fig. 3B). Cells were then washed three times with PBS and permeabilized in PBS supplemented with 0.5% (v/v) Triton X-100 for 10 min at room temperature. Cells were washed once with PBS, blocked with blocking buffer (0.1% Tween-20, 5% BSA, in PBS) for 1 h at room temperature, and labelled for 90 min with the following antibody dilutions: Fig. 3, rabbit anti-TOMM20 (1:200), and anti-Br(d)U (1:200); Fig. 4, mouse anti-TOM20 (1:200), anti-Br(d)U (1:200), anti-SUV3 (1:200), anti-GRSF1(1:200), anti-PNPT1 (1:200), and anti-Br(d)U (1:200) in blocking buffer. Cells were aspirated and washed three times for 5 min each in PBS and incubated with secondary antibodies in a blocking buffer at RT in the dilutions: anti-rabbit-AF488 (1,1,000) and anti-rat-AF555 (1:500) for Fig. 3. Finally, cells were aspirated, washed three times in PBS as before, stained with DAPI (Invitrogen, cat. #D1306) for 5 min, washed with PBS, and mounted using Prolong Glass Antifade Mountant (Thermo Fisher Scientific).

For EdU labelling of newly synthesized DNA, cells were treated as mentioned for BrU assays, but each glucose deprivation media was spiked with 20 µM EdU. Cells were treated for 24 h and then aspirated, washed twice in 37 °C PBS, and fixed as mentioned above. Cells were then permeabilized for 15 min and blocked for 30 min in solutions mentioned above. The blocking buffer was aspirated, and cells were washed once with PBS before treatment with the Click-iT reaction containing AF647-azide for 45 min. The reaction was aspirated, washed once with PBS, and incubated with primary antibodies in blocking buffer for 90 min at room temperature with the following dilutions: rabbit anti-TOMM20 (1:200) and anti-AF647 (1:200). The samples were then aspirated and washed three times in PBS for 5 min before secondary antibody incubation with the dilutions: anti-rabbit-AF488 (1:1,000) and anti-mouse-AF594 (1:1,000). Washing, DAPI staining, and mounting was carried out as above. Specific antibodies can also be found in Supplemental Table 1.

### Confocal Microscopy and Image Analysis

For BrU pulse-chase, EdU incorporation, degradosome, and J2 imaging experiments, Z-projection confocal images were acquired using a Leica TCS SP8 inverted microscope (Leica, Wetzlar, Germany) equipped with a white light laser covering 470-680 nm and a 405 nm laser line and spectral detection. Confocal images were taken using a 63x HC PL APO oil objective (Leica Germany NA1.4) sequentially between stacks with acquisition controlled by the LAS X (v. 3.5.2) software (Leica) and a logical size of 1024×1024. Pixel size for imaging was generally between 110-120 nm. For BrU and TOMM20 imaging the BrU sample was stained with Alexa Fleur 555 (AF555) and Alexa Fleur 488 (AF488), excited with the 553 or 499 laser lines, and detected from 563-640 or 511-545 nm respectively.

For degradosome (SUV3, GRSF1, and PNPT1), BrU, and TOMM20 imaging, staining was carried out using AF488, AF555, and Alexa Fleur 594 (AF594), and detection was carried out from 505-538, 556-575, and 607-666 nm respectively. These images were then deconvolved using the Leica Lightening deconvolution software on “adaptive” mode and a refractive index of 1.52. For dsRNA/J2 and TOMM20 imaging, the TOMM20 labelling and detection was carried out as for TOMM20/BrU imaging above while dsRNA/J2 was stained and detected AF594, excited with a 590 nm laser line, and detected from 607-676 nm. All DAPI signal was collected by excitation with the 405 nm laser line and detection between 431-468 nm. All image data was collected using hybrid detectors.

For live cell imaging on the same microscope, cells were maintained at 37 °C and 5% CO_2_. 2D images were acquired with simultaneous excitation at 488 nm and 645 nm, with detection between 493-630 nm and 652-776 nm, respectively. Images had a logical size of 1024×1024.

Images were initially saved in a .lif file. For analysis of BrU pulse-chase and EdU incorporation images, the maximum projection for each individual channel and an image containing DAPI and TOMM20 signals were exported as a .tiff from the Leica X software. Image segmentation and analysis were then completed in MATLAB (Mathworks). First, all channels were converted to grayscale for segmentation. The DAPI channel was then smoothed with a Gaussian filter and binarized to best cover the nucleus. Holes were then filled in the binary image, and the image was cleaned by removing objects with fewer than 25 pixels. The resulting binary DAPI channel was used as a mask over the original and grayscale BrU channel images to remove the nucleus. The overlayed DAPI and TOMM20 channel image was then used to pick out cells as individual ROIs using the drawfreehand function (Fig. S3A). The ROIs were used as masks over the nucleus-removed BrU image, and the intensity of each region was summed as the extranuclear BrU intensity of each cell per square micrometer. Similarly, TOMM20 intensity per µm^2^ was determined using each ROI over the original TOMM20 channel image.

For analysis of average MRG intensity from chase images, the nucleus-removed grayscale BrU image was first subjected to background subtraction of the bottom 8% intensity in a given image, followed by removal of small objects of less than 5 pixels and binarization. The binary image was then subject to watershed segmentation and removal of objects less than 5 pixels to yield a “cleaned” binary image. Object recognition was then carried out on the “cleaned” binary image and pixels of each object were extracted using the bwlabel and regionprops functions. Each object’s pixels were then extracted from the original BrU channel image and used to calculate the mean pixel intensity per MRG. Extranuclear dsRNA/J2 signal was measured from maximum intensity projections following nuclei removal is the same way as extranuclear BrU intensity measurements.

The analyses of degradosome components and 12 h BrU incorporation were carried out as follows: nuclei removal, extranuclear BrU, and TOMM20 intensity was measured as mentioned above from raw maximum projection images. Line-scan analysis was carried out on deconvolved maximum projection images by defining a line using the Matlab Bresenham line algorithm (Mathworks) using the bresenham() command and extracting profiles for each channel.

Live cell image analysis was completed by converting both channels to grayscale images. The sfGFP image contrast was then adjusted automatically using the imadjust function, and ROIs were drawn around each cell as previously mentioned using this image. The ROI was then masked over both miRFP670 and sfGFP original images and intensity was measured. ATP levels per cell were then assessed as the ratio of sfGFP to miRFP670 intensity per ROI.

### Western blotting

Cells treated in 6-well or 10 cm dishes were harvested by scraping, spun at 200 x *g* for 5 min, washed once with 5 ml ice-cold PBS, spun again, and finally lysed in a modified RIPA buffer (10 mM Tris-HCl pH 7.5, 150 mM NaCl, 1% Triton X-100, 0.5% sodium deoxycholate, 0.1% sodium dodecyl-sulfate, 10 mM nicotinamide, 330 nM trichostatin A, and protease inhibitor cocktail) for 30 min on ice. Lysates were spun at 17,709 x *g* for 20 min at 4 °C to remove cell debris, and the supernatant was collected for western blotting. Protein content was estimated using a BCA assay (Thermo Fisher Scientific), and 20-70 µg of total protein was used for each blotted sample. Samples were separated using 10% SDS-PAGE and transferred to a PVDF membrane. Blocking was performed with 5% dried milk in 1x TBST (1x TBS with 0.1% Tween-20) for 1 h at room temperature. Primary antibody incubation was performed overnight at 4 °C with antibodies and dilutions listed in Supplemental Table 1. After chemiluminescent detection on a Chemidoc Imaging System (BioRad), membranes were washed with water and stained in amido black staining solution (0.1% amido black, 40% ethanol, 10% glacial acetic acid), destained briefly in 5% acetic acid, and dried at room temperature for imaging total protein using a Chemidoc Imaging System. Band intensity was measured using ImageJ and normalized to a corresponding section of the membrane to the protein of interest using amido black intensity in each respective lane. Experimental samples were then compared to the average of the respective HG or DMSO control.

### Whole cell ATP content measurements

2,000 HEK293T cells were seeded into a 96-well dish and allowed to attach for 6-8 h. Medium was then aspirated, treatment medium was used to wash the cells once, and finally treatment medium was added to a final volume of 100 µl. The CellTiter-Glo® Luminescent Cell Viability Assay (Promega) reagent was then used according to the manufacturer’s specifications. Data are reported as ATP levels relative to the HG average value.

### Histone Extraction

Three 15 cm plates of HEK293T cells were used for each replicate of the three conditions: HG, NGBHB, and HGBHB mentioned above. Cells were resuspended and swelled in 3 ml of a hypo-lysis buffer (210 mM mannitol, 70 mM sucrose, 5 mM Tris-HCl pH 7.5, 1 mM EDTA) on ice for 10 min, then right before lysis a 2.5X homolysis buffer (525 mM mannitol, 175 mM sucrose,12.5 mM Tris-HCl pH 7.5, 2.5 mM EDTA, 25 mM NAM, 900 µM TSA, 1X PhosStop phosphatase inhibitor cocktail, EDTA-free 2.5X protease inhibitor cocktail, Roshe) was added before 50 strokes with a Dounce homogenizer. The lysis solution was then spun at 2505 RPM for 5 min to pellet nuclei. Nuclei pellets were then washed 1X with 1 ml ice cold PBS, and histones were extracted using 500 µl of 0.4N H_2_SO_4_ with rotation for 2 h at 4°C. Nuclear debris was then pelleted by spinning at 3400 x *g* for 10 min at 4 °C. The supernatant was collected and 125 µl of 100% trichloroacetic acid was added, and the mixture was incubated on ice for 1 h to precipitate proteins. The sample was then aspirated, and precipitants were washed twice with ice cold 100% acetone, each time with spinning at 3400 x *g* for 2 min and removing the supernatant. The sample was then allowed to air-dry at room temperature and dissolved in water. Sample purity was checked on a 12% SDS-PAGE gel with Coomassie staining, and protein concentration was determined with a BCA assay. 2 µg of total histone proteins from each sample were used for each lane in western blots for H3 and KBHB detection. Western blots were carried out as mentioned above in the “Western blotting” section.

### Statistical Analysis

Data were visualized using ggplot2 in R version 4.4.0. All T-tests and One-way ANOVAs with Tukey’s test were performed in R using the rstatix package (ns = nonsignificant, \**p*<0.05, \*\**p*<0.01, \*\*\**p*<0.001, \*\*\*\**p*<0.0001). T-tests were not corrected for multiple comparisons because our comparisons were pre-planned. Data collection and analysis were performed not blinded to experimental conditions.

## Author Contributions

Conceptualization, S.D.R and T.V.M.; methodology, S.D.R and J.C.; investigation, S.D.R., C.B., and S.C.C; formal analysis, S.D.R., C.B., and S.C.C; visualization, S.D.R.; writing – original draft, S.D.R and T.V.M.; writing – review and editing, S.D.R. and T.V.M.; fund acquisition, T.V.M.; supervision, T.V.M.

## Abbreviations

mtDNA: mitochondrial DNA
mtRNA: mitochondrial RNA
OXPHOS: oxidative phosphorylation
ISR: integrated stress response
BHB: β-hydroxybutyrate

## Acknowledgements

We would like to thank the members of the Debelouchina, Budin, Herzik, McHugh, Manor, and Zid labs at UC San Diego for sharing materials, cell lines, equipment, and expertise. We would like to thank Drs. Gerald S. Shadel, Brian M. Zid, Uri Manor, and L. Stirling Churchman (Harvard Medical School), as well as the Mishanina and Churchman labs for insightful discussion. RNA sequencing was conducted at the IGM Genomics Center, University of California, San Diego, La Jolla. We want to thank members of the UC San Diego School of Medicine Imaging Core: Jennifer Santini and Marcy Erb, as well as Carlos Alonso for guidance with confocal imaging. The UC San Diego School of Medicine Imaging Core is supported by the NINDS P30NS047101 grant.

---

**Supplemental Figure 1.**
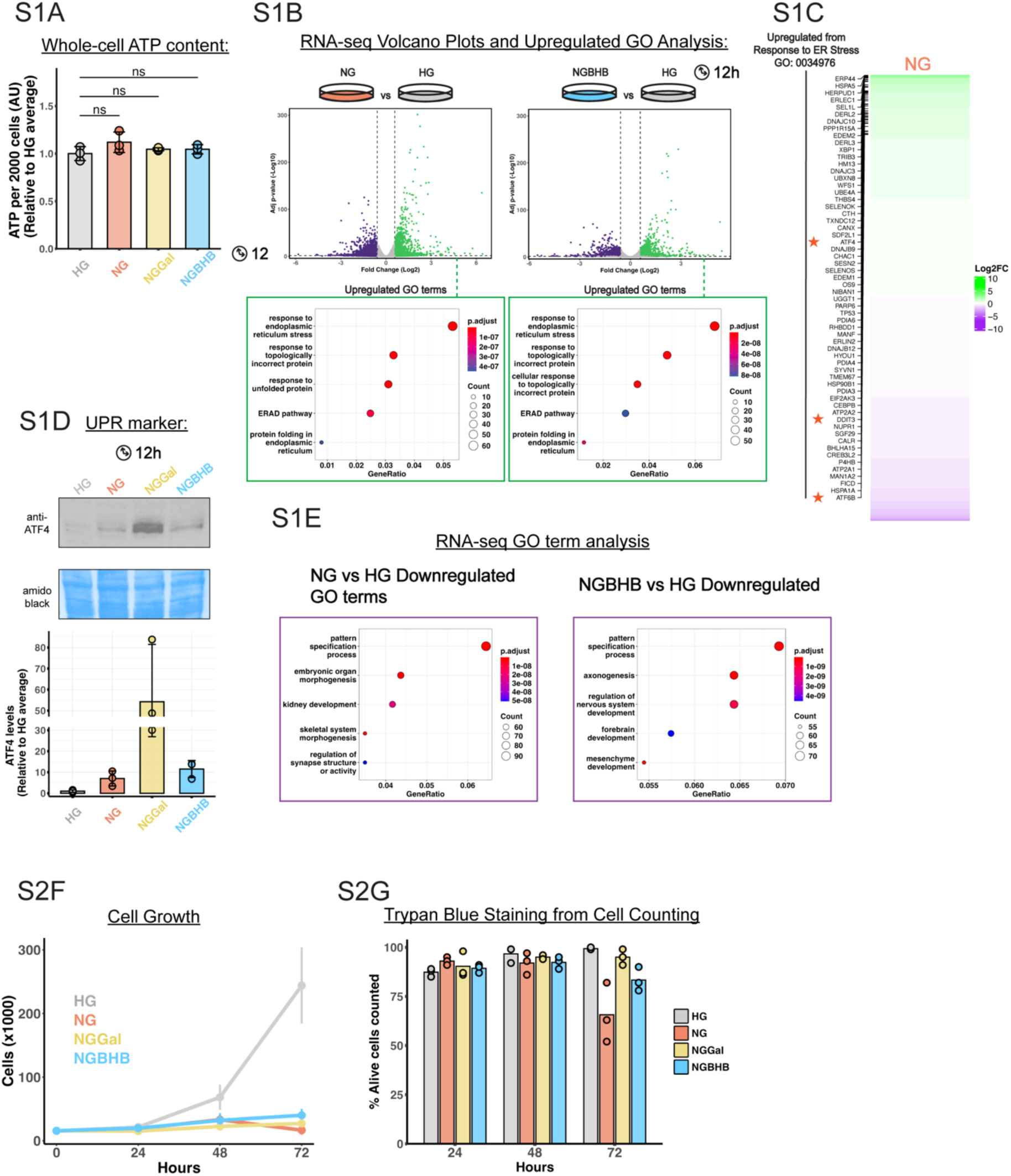
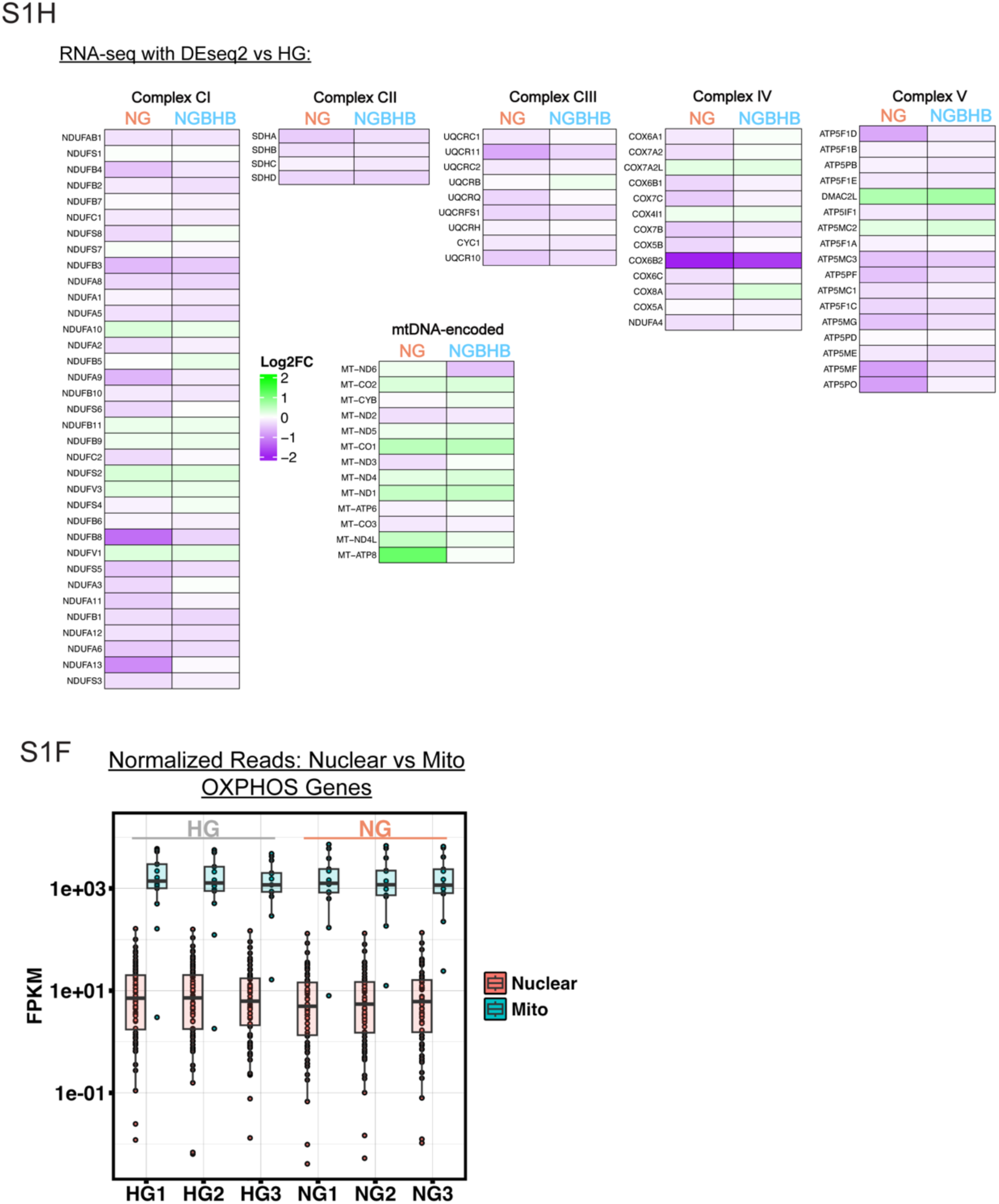
**A)** Whole-cell ATP content measurements with an ATP GLOW assay in cells deprived of glucose for 12 h (ns, non-significant; one-way ANOVA with Tukey’s multiple comparisons correction, data are means ± S.D.). **B)** RNA-seq volcano plots depicting differential gene expression analysis following 12 h treatments, with upregulated genes in green and downregulated genes in purple. Gene ontology (GO) analysis of upregulated genes from each dataset is shown in the bottom panels. Genes were considered “differentially expressed” if they showed fold changes >1.3 and adjusted p-value <0.05 (n = 3 biological replicates). **C)** Genes analyzed for gene expression analysis in NG vs HG with upregulated genes associated with the response to ER Stress (GO:0034976) shown to the left, and specific ISR transcription factors marked with orange stars. **D)** A representative western blot (top) showing ATF4 protein levels in glucose-deprived HEK293T cells, quantified (bottom) and normalized to high glucose (HG) average from 3 independent biological experiments. **E)** Gene ontology (GO) analysis of downregulated genes from transcriptomics comparing HEK293T cells 12 h treated NGGal and NGBHB compared to HG. (n = 3 biological replicates). **F)** Cell counting for HEK293T cells treated for 24, 48, and 72 hours in high glucose (HG), no glucose (NG), no-glucose-galactose (NGGal), and no-glucose-beta-hydroxybutyrate (NGBHB) (n = 3 biological replicates). **G)** Cell survival percentage determined by Trypan Blue staining from cells counted in F. **H)** Heatmap showing RNA-seq based log_2_ fold-change in expression of nuclear and mitochondrially encoded OXPHOS protein genes after a 12 h NG or NGBHB treatment, compared to HG (n = 3 biological replicates) **F)** RNA-seq analysis of nuclear and mitochondrially encoded OXPHOS protein genes as FPKM (n = 3 biological replicates).

**Supplemental Figure 2:**
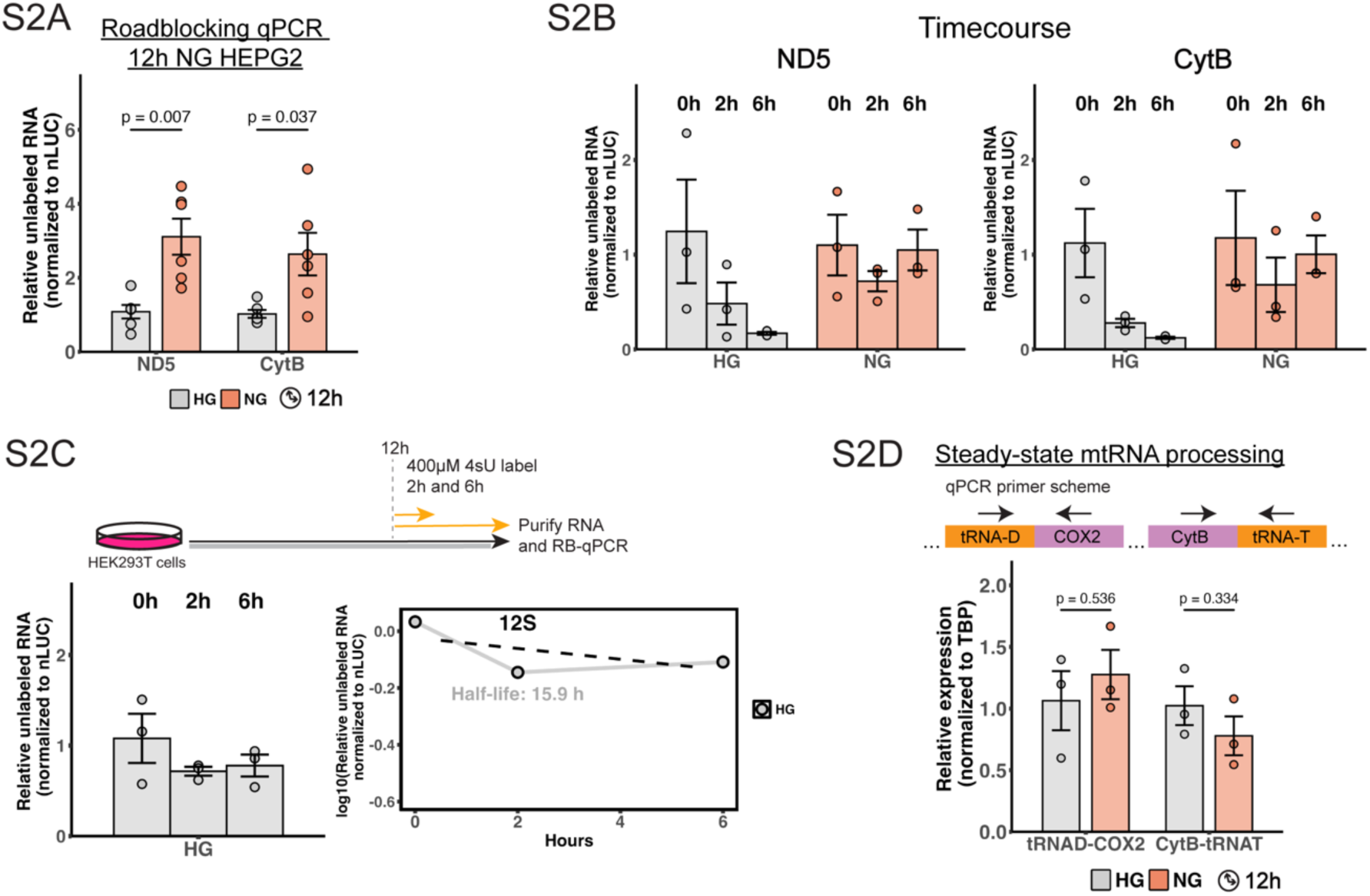
**A)** Roadblocking qPCR analysis for HEPG2 cells treated for 12 h in high glucose (HG) or no-glucose (NG) (n = 6 biological replicates, two-tailed Student’s *t*-test, data is mean ± s.e.m. with associated *P* values). **B)** Roadblocking qPCR analysis for ND5 and CytB in HEK293T cells treated with HG or NG for 12 h before 4sU incorporation for 0, 2, or 6 h (n = 3 biological replicates, normalized to nLUC RNA and the respective 0 h timepoint). **C)** Roadblocking qPCR analysis for the mitochondrial 12S ribosomal RNA (12S RNA) in HEK293T cells treated with HG for 12 h before 4sU incorporation for 0, 2, or 6 h (n = 3 biological replicates, normalized to nLUC RNA and the respective 0 h timepoint). **D)** Scheme and quantification for qPCR analysis of mitochondrial RNA processing in HEK293T cells treated in NG for 12 h compared to HG (n = 3 biological replicates, normalized to tata-box-binding protein (TBP), two-tailed Student’s *t*-test, data is mean ± s.e.m. with associated *P* values).

**Supplemental Figure 3:**
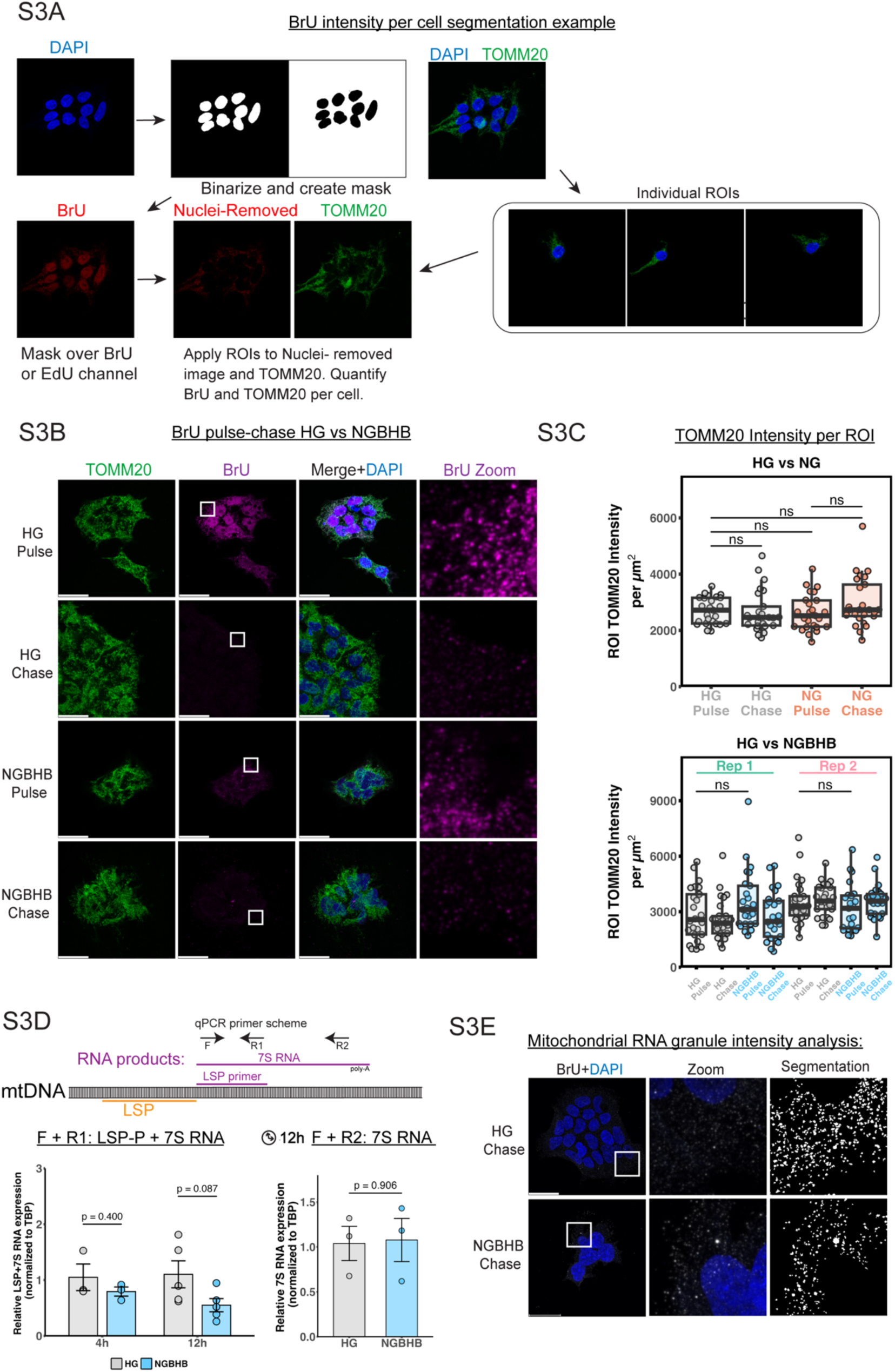
**A)** Example segmentation scheme for measurement of extranuclear BrU or EdU intensity in HEK 293T cells. This includes nucleus removal and ROI selection to represent individual cells. **B)** Representative images as maximum intensity projections for HG vs. NGBHB treated cells stained against TOMM20 (mitochondria) and BrU (newly synthesized RNA). Scale bar = 30 µm. **C)** Quantification of TOMM20 intensity per cell ROI for pulse and chase samples of HG vs NG (top), and HG vs. NGBHB (bottom, ns, non-significant; one-way ANOVA with Tukey’s multiple comparisons correction; n=50 cells per replicate, with 25 cells in each condition quantified, data represents individual cells and bars represent median, boxes represent the 1^st^ and 3^rd^ quartiles, and whiskers are no further than 1.5 times the interquartile range). **D)** Quantitative PCR (qPCR) primer scheme and analysis of the mitochondrial Light Strand Promoter primer region and 7S RNA in HEK293T cells treated for 12 h in HG or NGBHB (n = 3-6 biological replicates, normalized to tata-box-binding protein (TBP), two-tailed Student’s *t*-test, data is the mean ± s.e.m. and associated *p* value). **E)** Maximum intensity projection chase images for HG and NGBHB treated cells, and segmented mitochondrial RNA granules (left, MRGs) for quantification of average pixel intensity per MRG from samples in 3B (right).

**Supplemental Figure 4:**
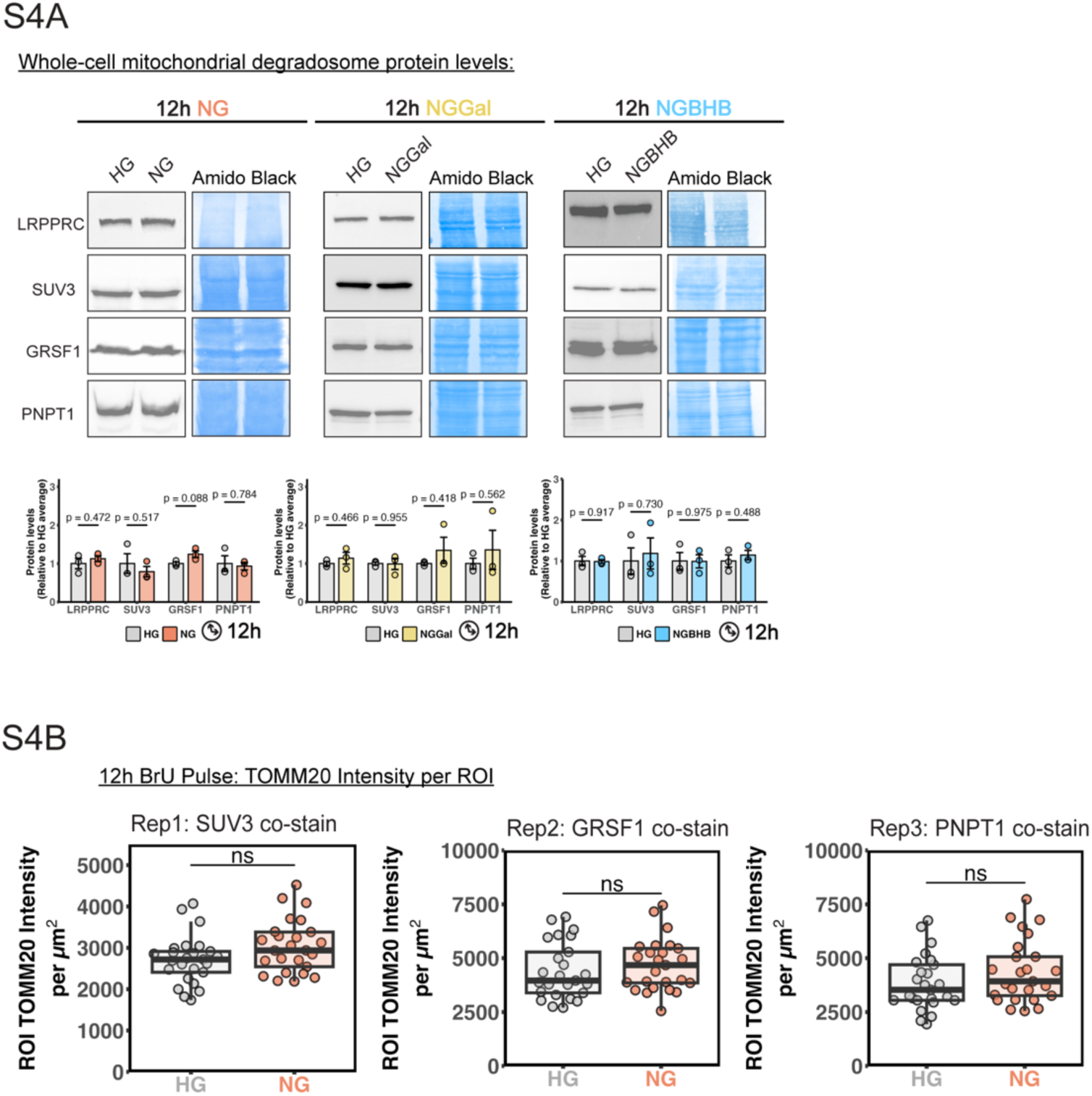
**A)** Quantification of TOMM20 intensity per cell ROI for 12 h NG pulse samples described in and chase samples of HG vs NG (top), and HG vs. NGBHB (ns, non-significant; two-tailed Student’s *t*-test; n=50 cells per replicate, with 25 cells in each condition quantified, data represents individual cells and bars represent median, boxes represent the 1^st^ and 3^rd^ quartiles, and whiskers are no further than 1.5 times the interquartile range).

**Supplemental Figure 5:**
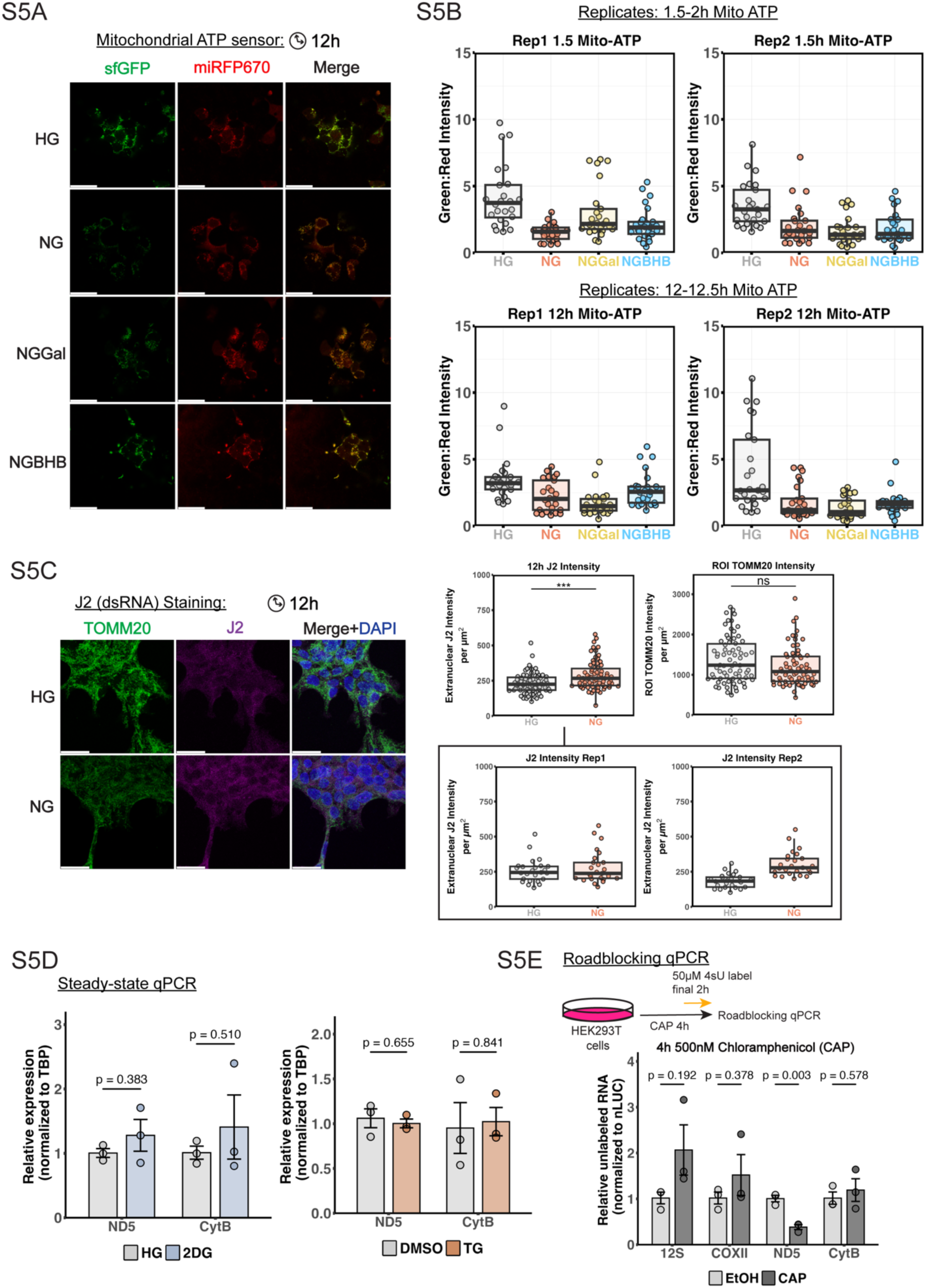
**A)** Representative 2D images of mitochondrial ATP sensor imaging in HEK293T cells during treatment for 12 h hours with glucose deprivation conditions. **B)** Replicate data for quantification of mitochondrial ATP per cell as the ratio of green:red intensity for samples treated in each condition for 1.5 and 12 h in Fig. 5B. Data represents individual cells and bars represent median, boxes represent the 1^st^ and 3^rd^ quartiles, and whiskers are no further than 1.5 times the interquartile range. **C)** Representative images of extranuclear dsRNA immunostaining with the J2 antibody of HEK293T cells treated with NG for 12 h. Cells were stained against TOMM20 (green, mitochondria) and J2 (magenta, dsRNA). **D)** Quantification of extranuclear dsRNA (J2) intensity and TOMM20 intensity per cell ROI from cells in S5C (top)(ns, non-significant; ***p<0.001, two-tailed Student’s *t*-test; n=60 cells per condition, with 30 cells in each condition quantified per replicate, data represents individual cells and bars represent median, boxes represent the 1^st^ and 3^rd^ quartiles, and whiskers are no further than 1.5 times the interquartile range). Replicate data for J2 measurements are boxed (below). **E)** Quantitative PCR (qPCR) analysis of representative mitochondrial transcripts shows steady pools of these RNAs after treatment with 25mM 2DG or 1µM thapsigargin (n = 3 biological replicates, normalized to tata-box-binding protein (TBP), data are means ± S.E.M. and associated *p* values). **F)** Roadblocking qPCR analysis for HEK293T cells treated for 12 h with 500nM chloramphenicol (n = 3 biological replicates, two-tailed Student’s *t*-test, data is mean ± s.e.m. with associated *P* values).

**Supplemental Figure 6:**
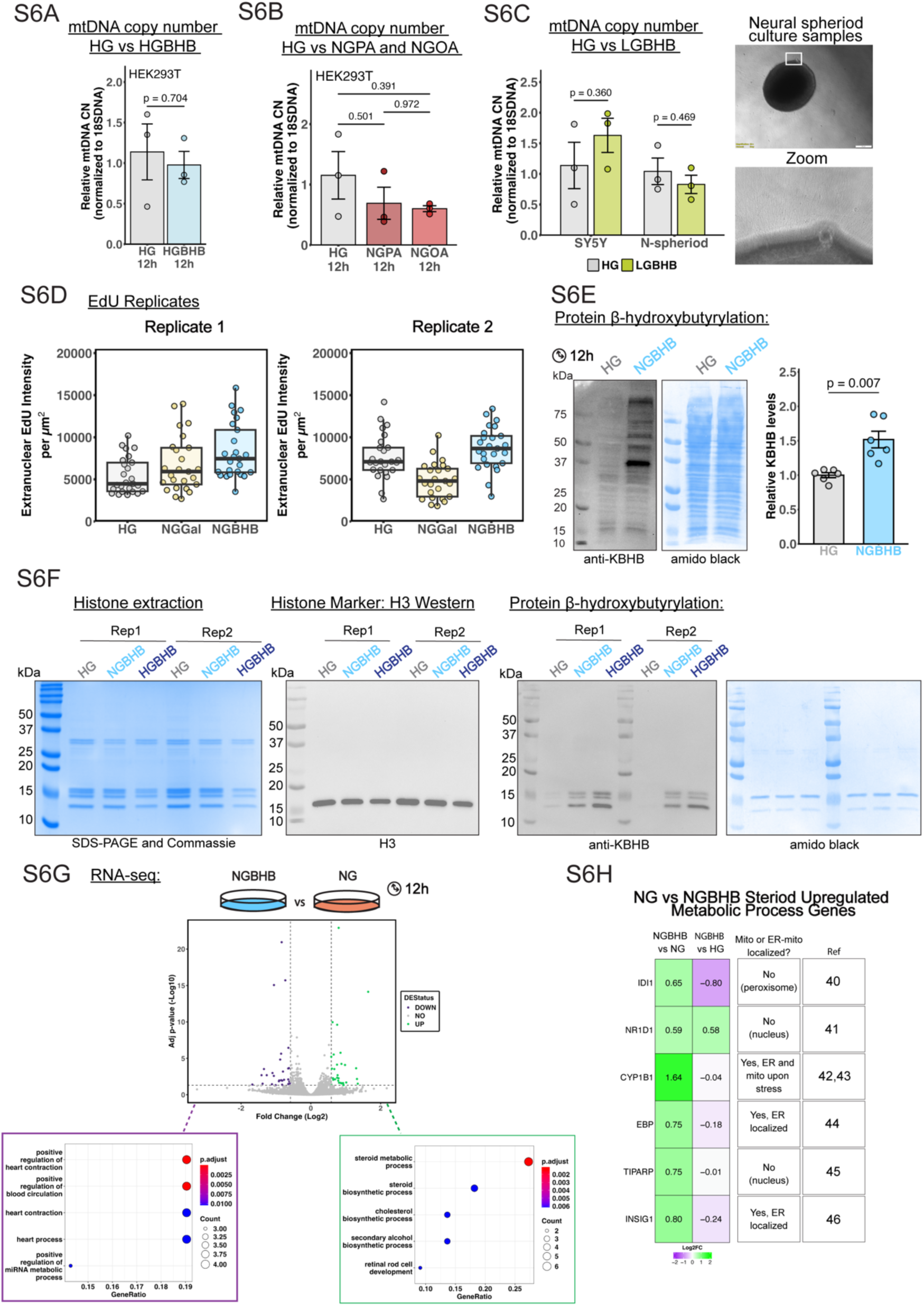
**A)** Quantitative PCR (qPCR) analysis of mitochondrial DNA copy number between 12 h high glucose (HG) and high-glucose with β-hydroxybutyrate (HGBHB) treated HEK293T. Mitochondrial DNA was detected with primers for CytB normalized to 18S DNA (n = 3 biological replicates, two-tailed Student’s *t*-test, data is mean ± s.e.m. with associated *p* values). **B)** Quantitative PCR (qPCR) analysis of mitochondrial DNA copy number between high glucose (HG) and no-glucose-palmitic acid (PA) and no-glucose-oleic-acid (OE) conditions in HEK293T cells treated for 12 h. mtDNA was detected as above (n = 3 biological replicates, two-tailed Student’s *t*-test, data is mean ± s.e.m. with associated *P* values). **C)** Quantitative PCR (qPCR) analysis of mitochondrial DNA copy number between high glucose (HG) and low-glucose-beta-hydroxybutyrate (LGBHB) conditions in SH-SY5Y cells and human neuronal spheroid culture treated for 12 h. (left, n = 3 biological replicates, two-tailed Student’s *t*-test, data is mean ± s.e.m. with associated *P* values) and images of human neuronal spheroid culture showing axons (right). **D)** Replicate data for the quantification of extranuclear EdU intensity per HEK293T cell in EdU pulse samples of high glucose (HG), no-glucose-galactose (NGGal), and no-glucose-BHB (NGBHB) treated cells (25 cells per condition and replicate, points represent individual cells, data represents individual cells and bars represent median, boxes represent the 1^st^ and 3^rd^ quartiles, and whiskers are no further than 1.5 times the interquartile range). **E)** Western blot analysis of lysine beta-hydroxybutyrylation (KBHB) from HEK293T cell lysate treated with HG or NGBHB for 12 h (top), and quantification of blots (below, n = 6 biological replicates, normalized to high glucose (HG) average value, two-tailed Student’s *t*-test with associated *P* values). **F)** SDS-PAGE gel for cell fractionation and histone extraction from HEK293T cells treated with HG, NGBHB, and HGBHB for 12 h (left), Histone H3 western blots of these samples (center-left), KBHB western blots of these samples (center-right), and amido black staining of the KBHB western blot (right). **G)** Volcano plots depicting differential gene expression analysis between HEK293T cells treated for 12 h in NGBHB or NG, with upregulated (green), downregulated genes (purple), and gene ontology (GO) analysis of upregulated genes from each dataset (bottom) shown. Genes were considered “differentially expressed” if they had fold changes >1.3 and adjusted p-value <0.05 (n = 3 biological replicates). **H)** Log_2_ fold-change data from differential gene expression analysis for upregulated genes related to the “steroid metabolic process” GO term in HEK293T cells from 12 h NGBHB vs 12 h NG treatment (n = 3 biological replicates).

**Supplemental Table 1.** Resources Table: Chemicals, kits, and antibodies.

| Reagents or Resource and use | Source | Identifier |
| --- | --- | --- |
| DMEM, High Glucose, glutamine, no pyruvate | Thermo Fisher Scientific | 11965092 |
| DMEM, No glucose, glutamine, no pyruvate | Thermo Fisher Scientific | 11966025 |
| DMEM/F12 Media | Thermo Fisher Scientific | 11320033 |
| DMEM, no glucose, no glutamine, no phenol red | Thermo Fisher Scientific | A1443001 |
| PBS, pH 7.4 | Thermo Fisher Scientific | 10-010-023 |
| Penicillin-Streptomycin-Glutamine | Thermo Fisher Scientific | 10378016 |
| Fetal Bovine Serum | Thermo Fisher Scientific | A5256701 |
| ATV solution | Thermo Fisher Scientific |  |
| GlutaMAX™ Supplement | Thermo Fisher Scientific | 35050061 |
| Sodium (R)-3-Hydroxybutanoate | Sigma-Aldrich | 298360 |
| D-glucose | Thermo Fisher Scientific | D16-500 |
| D-(+)-galactose | Thermo Scientific | A12813.30 |
| Mycoplasma PCR Detection Kit | Abcam | ab289834 |
| Trypan Blue Solution, 0.4% | Gibco | 15250061 |
| TRIZOL Reagent | Thermo Fisher Scientific | 15596026 |
| RNA ScreenTape and associated reagents | Agilent |  |
| Illumina Stranded mRNA Prep Kit | Illumina |  |
| NovaSeq X Plus Sequencing Reagents | Illumina |  |
| Seahorse XF RPMI Medium | Agilent | 103680-100 |
| Seahorse XF Cell Mito Stress Test Kit | Agilent | 103010-100 |
| PowerUp™ SYBR™ Green Master Mix | Thermo Fisher Scientific | A25741 |
| CellTiter-Glo® Luminescent Cell Viability Assay | Promega | G7570 |
| 5-Bromouridine | Sigma-Aldrich | 850187 |
| Click-iT™ EdU Imaging Kit | Thermo Fisher Scientific | C10086 |
| Poly-D-lysine | Thermo Fisher Scientific | A3890401 |
| ProLong™ Glass Antifade Mountant | Thermo Fisher Scientific | P36982 |
| Goat anti-Mouse-AF594 Antibody (IF, 1:1,000) | Abcam | ab150116 |
| Rat anti-Br(d)U Antibody (IF, 1:500) | Abcam | ab6326 |
| Goat anti-Rat-AF555 Antibody (IF, 1:1,000) | Abcam | ab150158 |
| Rabbit Anti-TOMM20 Antibody (IF, 1:200) | Abcam | ab186735 |
| Rabbit Anti-TOMM20 Antibody (IF, 1:200) | Proteintech | 66777-1-Ig |
| Goat anti-Rabbit-AF488 Antibody (IF, 1:1000) | Abcam | ab150077 |
| Rabbit Anti-mTFA Antibody (WB, 1:4,000-5,000) | Abcam | ab307302 |
| Mouse Anti-LRP130 (LRPPRC) Antibody (WB, 1:1,000) | Santa Cruz Biotechnology | sc-166178 |
| Rabbit Anti-SUPV3L1 Antibody (WB, 1:1,000, IF, 1:200) | Proteintech | 12826-1-AP |
| Rabbit Anti-TEFM Antibody (WB, 1:2,000) | Novus | NBP2-94080 |
| Rabbit Anti-GRSF1 Antibody (WB, 1:2,000, IF, 1:200) | Abcam | ab205531 |
| Rabbit Anti-PNPT1 Antibody (WB, 1:1,000, IF: 1:200) | Proteintech | 14487-I-AP |
| Rabbit Anti-ATF4 Antibody (WB, 1:1,000) | Cell Signaling Technology | #11815 |
| Mouse Anti-CHOP Antibody (WB, 1:1,000) | Cell Signaling Technology | #2895 |
| Mouse Double-stranded RNA (dsRNA) Antibody (J2) (IF, 1:200) | GenScript | A02181-40 |
| Rabbit Anti K-BHB Antibody (WB, 1:1,000) | PTMab | PTM-1201RM |
| Rabbit Anti H3 Antibody | Cell Signaling Technology | 9717S |
| Rabbit Anti-VDAC antibody (WB, 1:2,000) | Cell Signaling Technology | 4866T |
| Anti-Mouse IgG (H+L), HRP Conjugate (WB, 1:2,500) | Promega | W4021 |
| Anti-Rabbit IgG (H+L), HRP Conjugate (WB, 1:2,500) | Promega | W4011 |
| Mouse Anti-Alexa Fluor 647 Antibody (IF, 1:200) | IC Labs | M647-65A-400 |
| DAPI | Invitrogen | D1306 |
| RNA Clean & Concentrator™-5 | Zymo | R1013 |
| Quant-iT RNA Assay Kit | Thermo Fisher Scientific | Q33140 |
| GeneJET Genomic Purification Kit | Thermo scientific | K0721 |
| PureLink™ Genomic DNA Mini Kit | Invitrogen | K182001 |
| 4-thiouridine | Sigma-Aldrich | T4509 |
| Random Hexamer Primers | Thermo Scientific | SO142 |
| Q5 DNA Polymerase | New England Biolabs | M0491 |
| DpnI | New England Biolabs | R0176S |
| *E. coli* Poly(A) Polymerase | New England Biolabs | M0276S |
| DNA cleanup kit | New England Biolabs | T1130S |
| HiScribe® T7 Quick High Yield RNA Synthesis Kit | New England Biolabs | E2050S |
| SuperScript™ II Reverse Transcriptase | Invitrogen | 18064014 |
| Deoxy-D-Glucose | Fisher Scientific | AAL0733803 |
| Click-iT EdU Imaging Kit | Thermo Fisher Scientific | C10086 |
| Tween-20 |  |  |
| Pierce Rapid Gold BCA Assay Kit | Thermo Scientific | A53225 |
| Amido Black 10B | Thermo Fisher Scientific | BP124-10 |
| Nicotinamide | Thermo Scientific | 128271000 |
| Trichostatin A (TSA) | MedChemExpress | HY-15144 |
| ECL Western Blotting Substrate | Promega | W1015 |
| Thapsigargin | Cell Signaling Technology | 12758S |
| PhosSTOP | Roche | 4906845001 |

**Supplemental Table 2:**
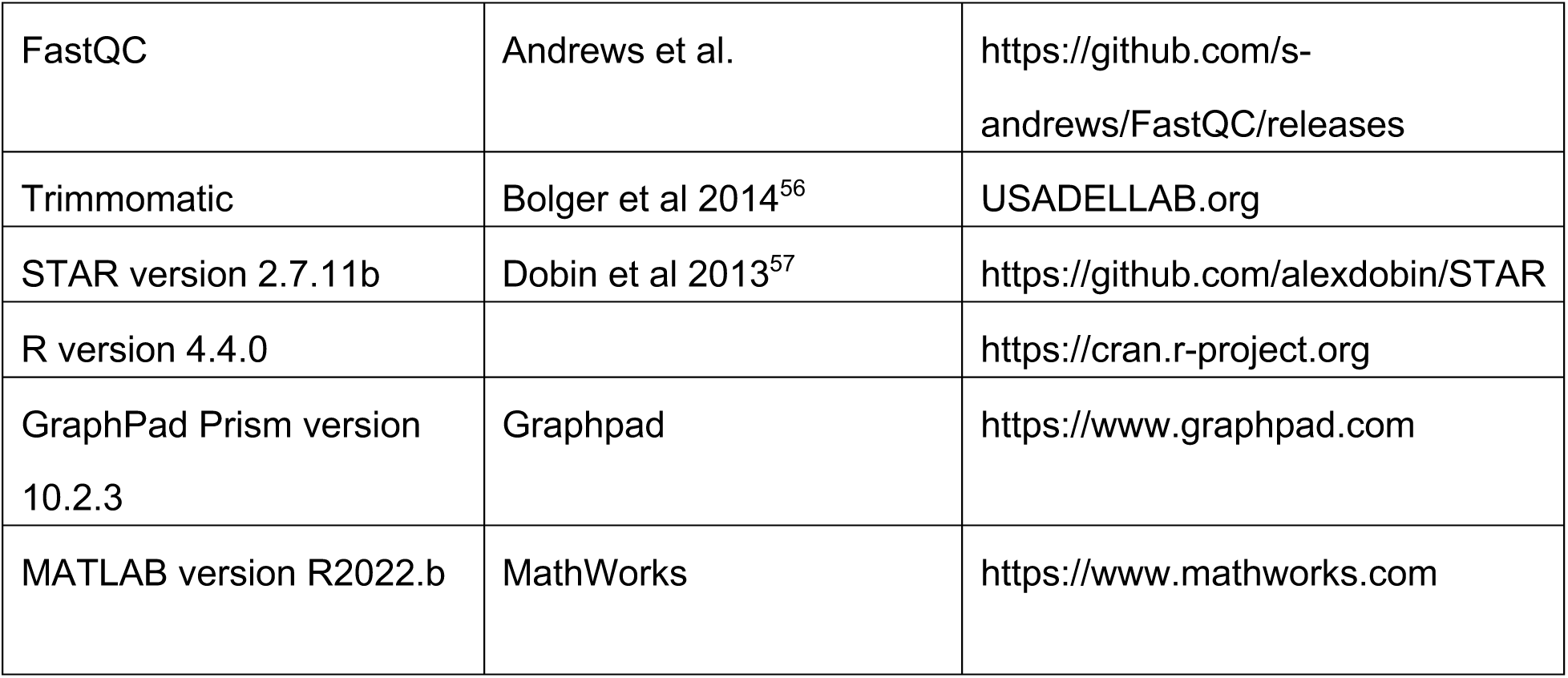
Software.

**Supplemental Table 3:** Plasmids.

| Plasmid | Source | Identifier |
| --- | --- | --- |
| nLUC-containing plasmid | Gift from Zid Lab (available on Addgene) | Addgene 199755 |
| pAAV.CAG.(mito).iATPSnFR2.A95A.A119L.miRFP670nano3 | Addgene | 209723 |

**Supplemental Table 4:**
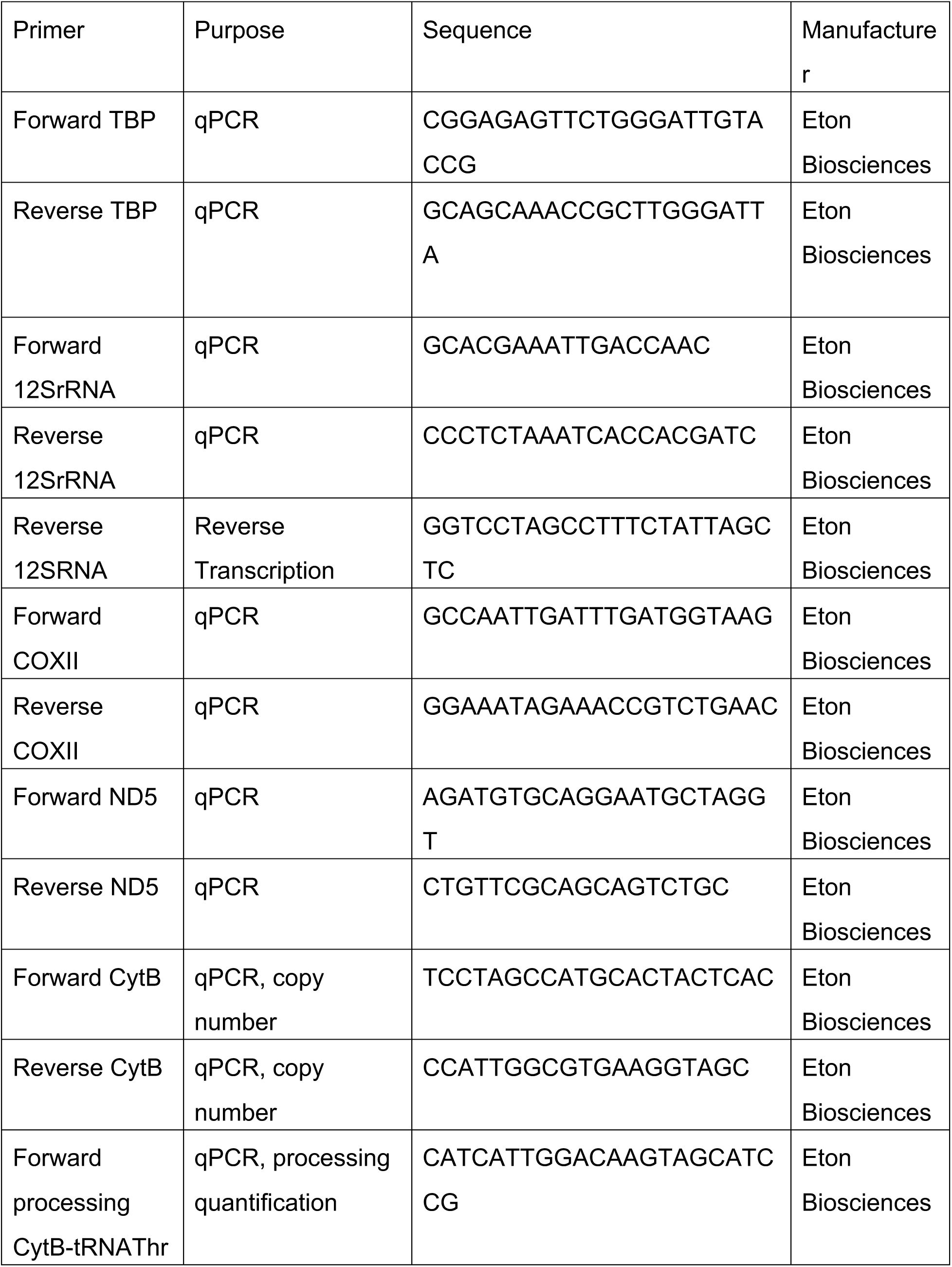
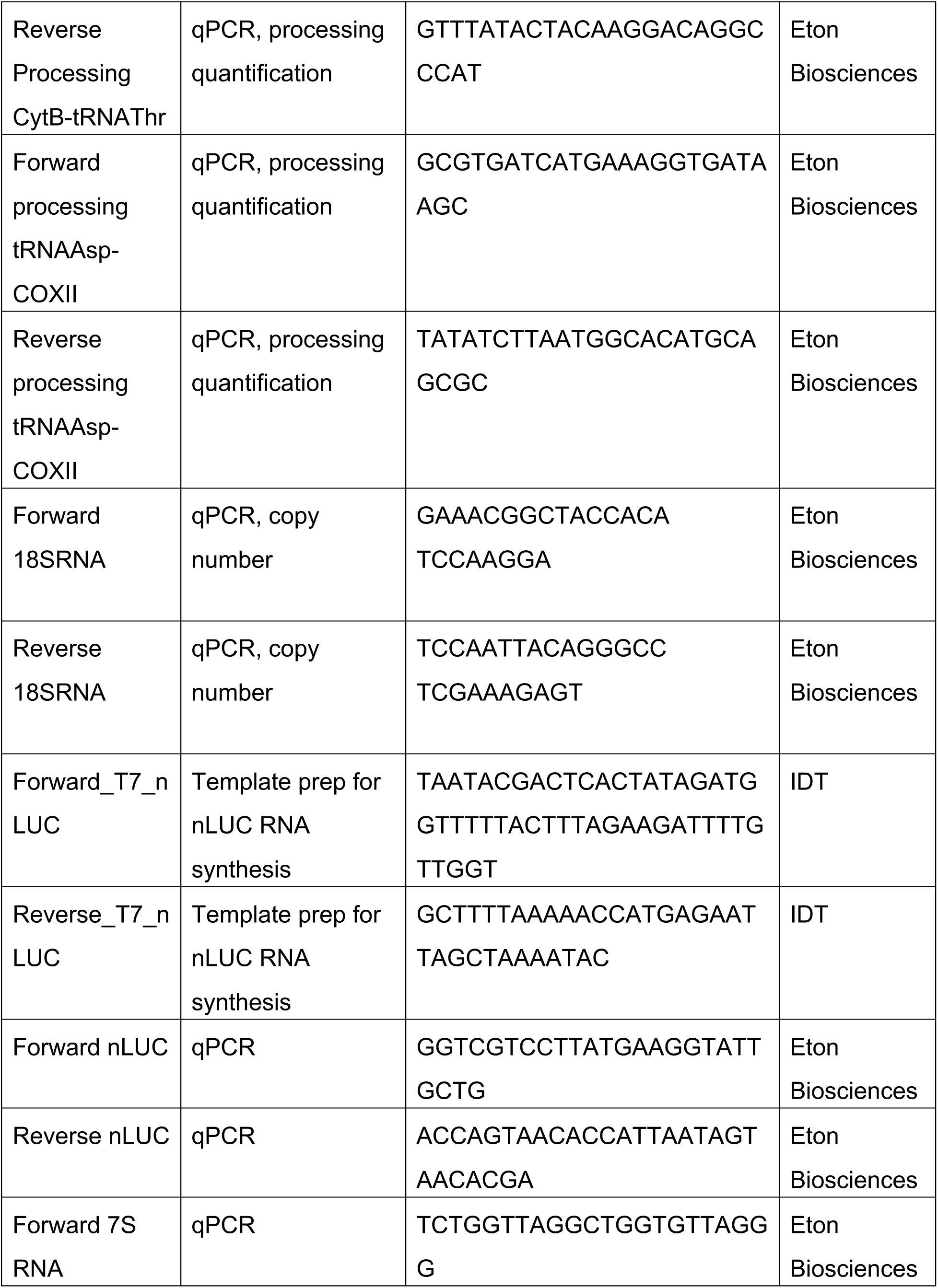
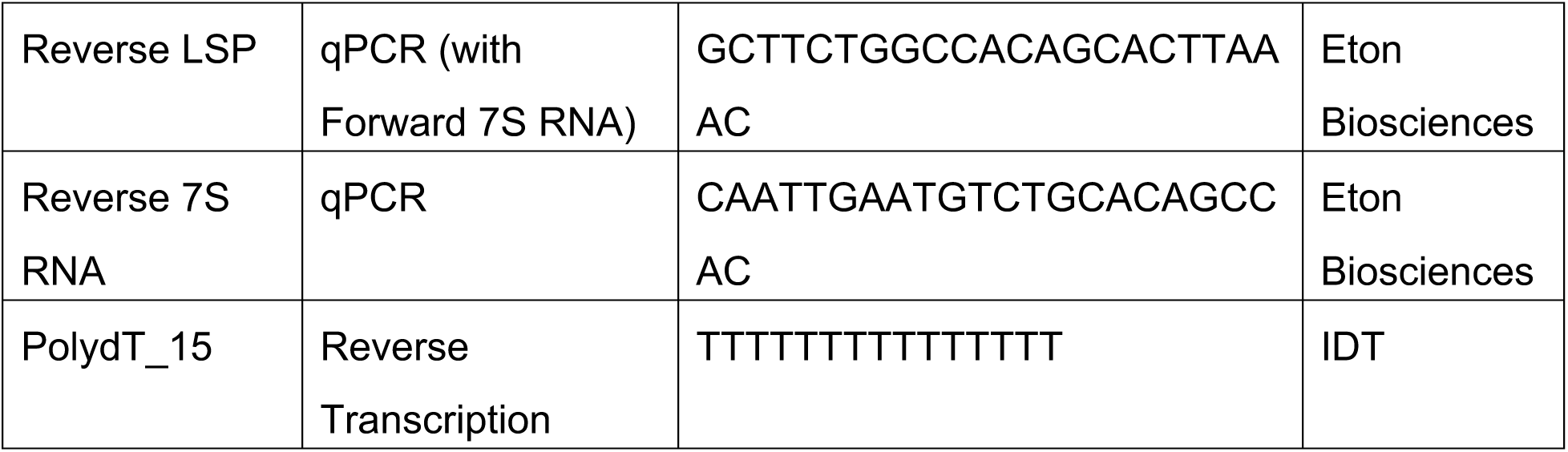
Oligonucleotides.

